# Type III interferons induce pyroptosis in gut epithelial cells and delay tissue restitution upon acute intestinal injury

**DOI:** 10.1101/2022.03.04.482997

**Authors:** Benedetta Sposito, Julien Mambu, Katlynn Bugda Gwilt, Lionel Spinelli, Natalia Andreeva, Franck Galland, Philippe Naquet, Vanessa Mitsialis, Jay R Thiagarajah, Scott B Snapper, Achille Broggi, Ivan Zanoni

## Abstract

Tissue damage and repair are hallmarks of the inflammatory process. Despite a wealth of information focused on the mechanisms that govern tissue damage, mechanistic insight on how inflammatory immune mediators affect the restitution phase is lacking. Here, we investigated how interferons influence tissue restitution after damage of the intestinal mucosa driven by inflammatory or physical injury. We found that type III, but not type I, interferons serve a central role in the restitution process. Type III interferons induce the upregulation of ZBP1, caspase activation, and cleavage of gasdermin C, and drive epithelial cell death by pyroptosis, thus delaying tissue restitution. We also found that this pathway is transcriptionally regulated in IBD patients. Our findings highlight a new molecular signaling cascade initiated by the immune system that affects the outcome of the immune response by delaying tissue repair and that may have important implications for human inflammatory disorders.

## Introduction

The immune system evolved to protect the host from external or internal threats, as well as to maintain homeostasis of the organs and tissues. The strong interrelationship between these two functions of the immune system is best exemplified during the restitution phase that follows mucosal damage, occurring as a consequence of an immune response. The skin, the lungs, the gut, and other mucosae are constantly exposed to microbial and/or physical perturbations and harbor multiple immune and non-immune cells that sense the presence of hostile environmental or endogenous factors and mount a defensive response. The causative agent of this response, the response itself, or both, may lead to tissue damage. Tissue damage sensing by tissue-resident as well as newly recruited cells initiates a complex cascade of cellular and molecular processes to restore tissue functionality and homeostasis, or to adapt to persistent perturbations (Meizlish, Franklin et al. 2021).

The gastrointestinal tract represents an ideal tissue to explore the mechanisms underlying the exquisite balance between tissue damage and repair orchestrated by the immune system. In the intestine, immune cells, epithelial cells, and commensal microbes are in a dynamic equilibrium. A monolayer of highly specialized epithelial cells separates the gut lumen from the underlying lamina propria. The interplay between microbiota-derived inflammatory cues and the host cells in the intestine profoundly impacts the biology of the gut, both during homeostasis, inflammation, and damage responses. The lamina propria hosts a large variety of immune and non-immune cells that detect alterations in the functioning as well as in the integrity of the epithelial barrier and mount an immune response. The fine equilibrium between the microbiota, the epithelial barrier, and the immune system is lost during inflammatory bowel diseases (IBDs). IBDs are a group of heterogeneous diseases, whose pathogenesis is associated with genetic and environmental factors, that are characterized by a dysregulated immune response (Danese and Fiocchi 2011, Roda, Chien Ng et al. 2020). Along with a heightened inflammatory response, IBDs are characterized by the breach of the intestinal barrier and a defective repair response that compromises mucosal homeostasis. Therefore, the ability of immune mediators to influence epithelial repair has an important impact on the pathogenesis of IBDs. Indeed, the promotion of mucosal healing has been recognized as a major therapeutic challenge for the management of IBDs (Pineton de Chambrun, Peyrin-Biroulet et al. 2010).

We previously showed that a group of interferons (IFNs), known as type III IFNs or IFN-λ (Kotenko, Gallagher et al. 2003, Sheppard, Kindsvogel et al. 2003, Prokunina-Olsson, Muchmore et al. 2013), limits inflammation in a mouse model of colitis by dampening the tissue-damaging functions of neutrophils (Broggi, Tan et al. 2017). IFN-λ, as type I IFNs, plays potent anti-microbial roles, but, in contrast to type I IFNs, also preserves gut functionality by limiting excessive damage (Broggi, Granucci et al. 2020). The limited damage is largely explained by the fact that the expression of the IFN-λ receptor (IFNLR) is mainly restricted to epithelial cells and neutrophils. In contrast, type I IFNs act systemically and play potent inflammatory activities on immune and non-immune cells thanks to the broad expression of the type I IFN receptor (IFNAR). The local activity of IFN-λ at mucosal tissues, thus, limits the extent of activation of immune cells, preventing excessive tissue damage, while preserving the anti-microbial functions of IFN-λ (Broggi, Granucci et al. 2020).

Although we and others have shown that IFN-λ limits intestinal tissue damage, the involvement of this group of IFNs during tissue restitution of the gut is more controversial. Indeed, IFN-λ and type I IFNs have been shown to function in a balanced and compartmentalized way to favor re-epithelization by acting, respectively, on epithelial cells or immune cells resident in the lamina propria (McElrath, Espinosa et al. 2021). IFN-λ has been also proposed to facilitate the proliferation of intestinal epithelial cells via STAT1 signaling (Chiriac, Buchen et al. 2017) and to partially enhance gut mucosal integrity during graft versus host disease (Henden, Koyama et al. 2021). On the other hand, though, IFN-λ and/or the IFNLR were found to be upregulated in IBD patients (Chiriac, Buchen et al. 2017, Gunther, Ruder et al. 2019). Also, systemic and prolonged overexpression of IFN-λ in mice favored the death of Paneth cells, a group of cells that can facilitate epithelial cell regeneration by acquiring stem-like features (Schmitt, Schewe et al. 2018), and by regulating the balance of epithelial growth factors in the stem cell niche (Sato, van Es et al. 2011).

In keeping with a possible detrimental role of IFN-λ during an inflammatory response at mucosal surfaces, we and others have recently demonstrated that IFN-λ delays the proliferation of lung epithelial cells in murine models of persistent viral infections (Broggi, Ghosh et al. 2020, Major, Crotta et al. 2020). Also, that IFN-λ production in the lower respiratory tract of COVID-19 patients is associated with increased apoptotic and decreased proliferative transcriptional programs, and characterizes SARS-CoV-2-infected individuals with severe-to-critical outcomes (Sposito, Broggi et al. 2021). Whether IFN-λ plays similar roles in the intestine, and the molecular mechanisms initiated by this group of IFNs to exert their functions during gut restitution, remain unknown.

Here, by exploiting conditional knock-out mice that do not respond to IFN-λ only in intestinal epithelial cells or in neutrophils, *ex vivo* transcriptomics, and biochemical assays, as well as intestinal organoids *in vitro*, we dissected the role of IFN-λ during tissue repair secondary to either an inflammatory insult or to radiation damage. Our data reveal a new molecular cascade initiated by IFN-λ that culminates in the activation of ZBP1 and of gasdermin C (GSDMC), in the induction of pyroptosis and results in delayed gut restitution.

## Results

### IFN-λ delays tissue repair of the inflamed gut

We previously showed that, in the acute inflammatory phase of the dextran sulfate sodium (DSS) model of colitis, IFN-λ signaling in neutrophils dampens reactive oxygen species production and neutrophil degranulation, and thus restrains intestinal damage (Broggi, Tan et al. 2017). To assess the involvement of IFN-λ during the restitution phase of the DSS colitis model, we injected, or not, recombinant (r)IFN-λ in mice after DSS-induced inflammation has peaked. We confirmed that rIFN-λ administration upregulated interferon-stimulated genes (ISGs) in the colon of DSS-treated mice (**Figure S1A**). Mice administered rIFN-λ, but not vehicle controls, showed persistent weight loss, reduced colon length, and prolonged tissue damage as measured by histology (**Figure 1A-C**). These data suggest that IFN-λ delays tissue restitution in mice encountering colitis.

**FIGURE 1.**
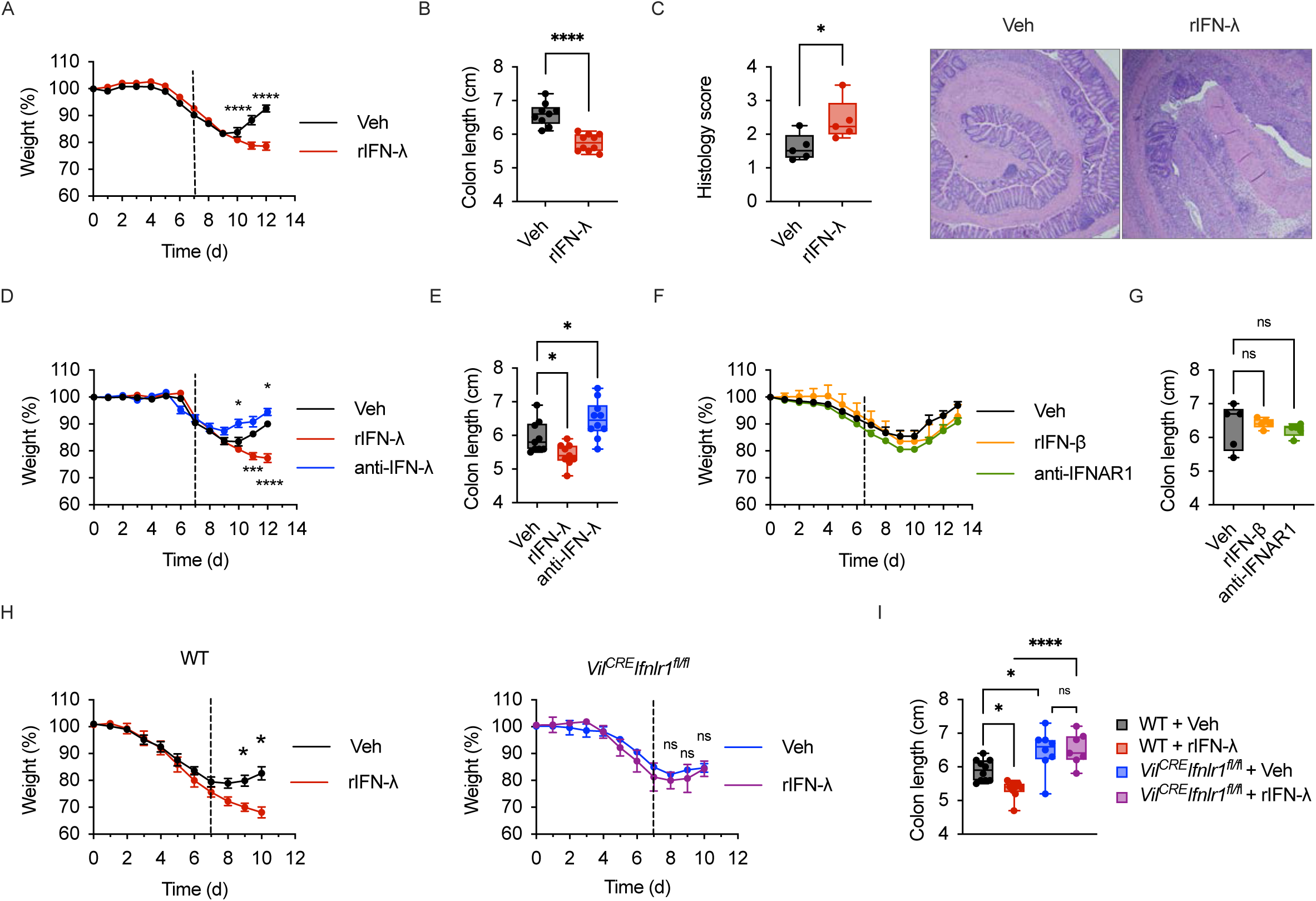
IFN-λ inhibits tissue recovery after DSS-colitis. (**A-C**) WT mice were treated with 2.5% DSS for 7 days (dotted line indicates the end of DSS administration). Upon DSS withdrawal on Day 7, mice were injected intraperitoneally (i.p.) with 50 µg kg^-1^day^-1^ of rIFN-λ for five consecutive days. Weight (**A**), colon length (**B**), histological score and representative histology images (**C**) are depicted. (**D, E**) WT mice were treated with DSS for 7 days as in (**A-C**). Upon DSS withdrawal mice were injected i.p. with either 50 µg kg-1day-1 of rIFN-λ or 12.5 mg kg^-1^day^-1^ of anti-IFN-λ2,3 antibody. Weight (**D**), and colon length (**E**) are depicted. (**F, G**) Mice were treated with DSS as in (**A-C**). Upon DSS withdrawal mice were injected i.p. with 50 mg kg^-1^day^-1^ of rIFN- β or 12.5 mg kg^-1^day^-1^ of anti-IFNAR1 antibody. Weight (**F**), and colon length (**G**) are depicted. (**H, I**) WT (**H**, *left panel*) or *Vil^CRE^Ifnlr1*^fl/fl^ mice (**H**, *right panel*) were treated with 2.5% DSS for seven days (dotted line). Upon DSS withdrawal mice were injected i.p. with 50 µg kg^-1^day^-1^ of rIFN-λ. Weight (**H**), and colon length (**I**) are depicted. (**A, D, F, H**) Mean and SEM of 5 mice per group are depicted. Two-way ANOVA with Tukey correction for multiple comparisons was utilized. (**B, C, E, G, I**) Box plots are depicted. Each dot represents a mouse. Median, range and interquartile range are depicted. Statistics: (**B, C**) Unpaired t test. (**E, G**) One-way ANOVA with Dunnett correction for multiple comparisons. (**I**) Two-way ANOVA with Šidak correction for multiple comparisons. Data representative of 3 independent experiments. ns= not significant (p > 0.05); *p < 0.05;**p < 0.01; ***p < 0.001; ****p < 0.0001.

Next, we tested whether the endogenous IFN-λ, which is produced during colitis development (Broggi, Tan et al. 2017), also affects the restitution phase. After the peak of the inflammatory process induced by DSS administration, mice were treated with a blocking antibody directed against IFN-λ and compared to mice treated with DSS, in the presence or absence of rIFN-λ. Our data demonstrated that inhibition of endogenous IFN-λ facilitates tissue restitution as measured by increased weight gain and colon lengthening (**Figure 1D, E**). Notably, we found that ISG levels in epithelial cells were significantly decreased in mice treated with the anti-IFN-λ antibody (**Figure S1A**), suggesting that IFN-λ, rather than type I IFNs, plays a major role in driving gene transcription during the repair phase of colitis. To directly test the involvement of type I IFNs in the restitution phase of DSS-induced colitis, we either blocked type I IFN signaling using an anti-IFNAR antibody, or added rIFNβ, 7 days after DSS administration. In keeping with a key role of IFN-λ in regulating mucosal epithelial responses, none of the treatments aimed at targeting type I IFNs affected tissue repair (**Figure 1F, G**). Accordingly, ISG levels in colonocytes were not altered under these experimental conditions compared to control mice (**Figure S1B**).

While intestinal epithelial cells are the major effector cell type during mucosal restitution, other cells, including immune cells, can participate in modulating tissue repair. Since intestinal epithelial cells and neutrophils are the two cell types that respond to IFN-λ in the gut of mice (Broggi, Tan et al. 2017), we used conditional knock out mice that do not express the IFNLR either in intestinal epithelial cells (*Vil^CRE^Ifnlr1^fl/fl^* mice) or neutrophils (*Mrp8^CRE^Ifnlr1^fl/fl^* mice). *Ifnlr1^fl/fl^* (WT) littermates were used as controls. In contrast to WT littermates, administration of rIFN-λ to *Vil^CRE^Ifnlr1^fl/fl^* mice did not delay tissue restitution as measured by weight change (**Figure 1H**). *Vil^CRE^Ifnlr1^fl/fl^*mice in which intestinal epithelial cells do not respond to IFN-λ showed a faster recovery as measured by a significant increase in colon length, regardless of the presence or absence of rIFN-λ (**Figure 1I**). In contrast to *Vil^CRE^Ifnlr1^fl/fl^* mice, *Mrp8^CRE^Ifnlr1^fl/fl^* behaved similarly to their WT counterpart, in the presence or absence of rIFN-λ (**Figure S1C**). These data demonstrate that, in contrast to the acute inflammatory phase of colitis, epithelial cells, not neutrophils, are the major responders to endogenous, as well as exogenous, IFN-λ and that IFN- λ signaling in epithelial cells delays tissue restitution.

### IFN-λ delays the tissue restitution phase that follows radiation damage

Repair of the gut epithelial monolayer is a complex process, and the regenerative capacity of intestinal stem cells (ISCs) plays a critical role (Blanpain and Fuchs 2014). To target ISCs and assess the direct involvement of IFN-λ during gut restitution, we employed a well-characterized model of epithelial damage resulting from exposure to ionizing radiations (Kim, Yang et al. 2017). In this model, radiation induces widespread epithelial cell death in the small intestine, with a particularly dramatic effect on cycling ISCs that reside at the bottom of the small intestinal crypt. Cell death is followed by repair of the damaged epithelial crypts and return to homeostasis. Three to four days after radiation injury, during the peak of the repair response, crypt regeneration was assessed in WT mice, WT mice administered exogenous rIFN-λ, or *Ifnlr1*^-/-^ mice. We found that mice that received rIFN-λ showed reduced regeneration of the crypts, while *Ifnlr1*^-/-^ mice had an increased number of crypts (**Figure 2A**). Notably, in the small intestine of irradiated mice, similarly to what we observed in the colon of mice exposed to DSS, endogenous IFN-λ, but not type I IFN, signaling caused the delay in tissue restitution (**Figure 2B**). Similarly, ISG induction in epithelial cells was dependent on IFN-λ, rather than type I IFNs (**Figure S2A**).

**FIGURE 2.**
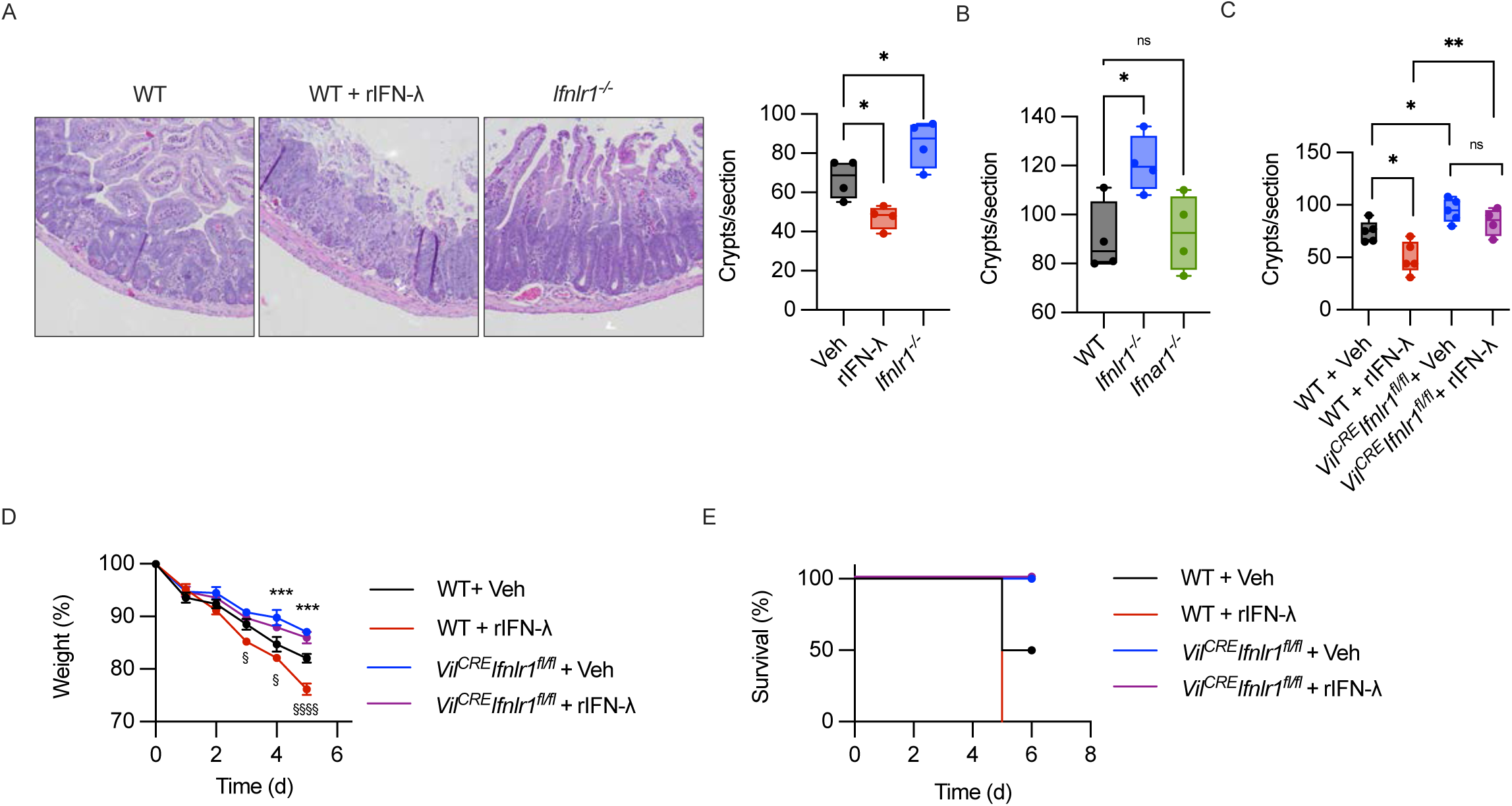
IFN-λ impairs epithelial regeneration after radiation damage. (**A**) WT mice and *Ifnlr1^-/-^*received 11 Gy ionizing radiation, with lead shielding of the upper body. WT mice were either administered 50 mg kg^-1^day^-1^ of rIFN-λ (WT + rIFN-λ), or the same volume of saline vehicle (WT + Veh). Tissue repair in the small intestine was evaluated 96h after irradiation by counting the number of intact crypts per histological section (Crypts/section). *Left panel*: Representative histological images; *right panel*: quantification. (**B**) WT, *Ifnlr1^-/-^*, or *Ifnar1^-/-^* mice were irradiated as in (**A**). Quantification of intact crypts per histological section is depicted. (**C**) *Vil^CRE^ Ifnlr1*^fl/fl^ mice or WT mice were irradiated as in (**A**) and treated with either 50 mg kg^-1^day^-1^ of rIFN-λ (rIFN-λ), or saline vehicle (Veh). Quantification of intact crypts per histological section is depicted. (**D, E**) *Vil^CRE^ Ifnlr1*^fl/fl^ mice or WT mice were treated with 14 Gy of ionizing radiation and treated with either 50 mg kg^-1^day^-1^ of rIFN-λ (rIFN-λ), or saline vehicle (Veh) and followed over time. Weight (**D**) and survival (**E**) are depicted. Statistical comparison between “WT+ Veh” and “*Vil^CRE^Ifnlr1^fl/fl^*+ Veh” are depicted as (*), comparison between “WT+ Veh” and “WT + rIFN-λ” is depicted as (§). (**A-C**) Box plots are depicted. Each dot represents a mouse. Median, range and interquartile range are depicted. (**D**) Mean and SEM are depicted. Statistics: (**A, B**) One-way ANOVA with Dunnett correction for multiple comparisons. (**C**) Two-way ANOVA with Šidak correction for multiple comparison. (**D**) Two-way ANOVA with Tukey correction for multiple comparisons. ns= not significant (p > 0.05); *p < 0.05;**p < 0.01; ***p < 0.001; ****p < 0.0001.

Next, we assessed the nature of the cell types that respond to IFN-λ in the irradiated small intestine. When *Vil^CRE^Ifnlr1^fl/fl^* mice and WT littermates were used, we found that the number of crypts three days post-radiation was significantly increased in *Vil^CRE^Ifnlr1^fl/fl^* mice compared to WT mice (**Figure 2C, S2B**). We also demonstrated that exogenous rIFN-λ does not affect the number of crypts in *Vil^CRE^Ifnlr1^fl/fl^* mice, while delaying tissue restitution in WT littermates (**Figure 2C, S2B**). In keeping with a key role for epithelial cells, but not neutrophils, in responding to IFN-λ during tissue restitution, we found that *Mrp8^CRE^Ifnlr1^fl/fl^* mice didn’t show significant differences compared to their WT littermates (**Figure S2C**). No differences were measured in the number of crypts of non-irradiated mice regardless of their capacity to respond, or not, to IFN-λ (**Figure S2D**). Finally, we followed over time irradiated *Vil^CRE^Ifnlr1^fl/fl^*mice or WT littermates, treated or not with rIFN-λ. We found that WT mice irradiated and treated with rIFN-λ lost significantly more weight than irradiated WT mice, and all died (**Figure 2D, E**). Notably, WT littermates lost significantly more weight compared to *Vil^CRE^Ifnlr1^fl/fl^* mice, treated or not with rIFN-λ (**Figure 2D**). In contrast, *Vil^CRE^Ifnlr1^fl/fl^*mice treated or not with rIFN-λ showed a very similar behavior (**Figure 2D, E**). Overall, these data demonstrate that epithelial cell regeneration and tissue restitution in the small intestine of irradiated mice is inhibited in the presence of IFN-λ. Also, that IFN-λ delays repair by acting on intestinal epithelial cells.

### IFN-λ dampens regenerative and proliferative transcriptional programs in intestinal epithelial cells

To determine the transcriptional programs initiated by IFN-λ to delay tissue restitution, we isolated intestinal crypts from the small intestine of *Vil^CRE^Ifnlr1^fl/fl^* mice or WT littermates that have been irradiated and performed targeted transcriptomics analysis (RNAseq). In keeping with a major role of IFN-λ-dependent responses in the intestine, when we performed gene ontology (GO) enrichment analyses, IFN-signaling related pathways, as well as anti-viral or anti-bacterial pathways, were highly enriched in WT epithelial cells, compared to *Vil^CRE^Ifnlr1^fl/fl^* (**Figure 3A**). In contrast, GO terms associated with cell migration and extracellular remodeling, which are linked to higher efficiency in the closure of mucosal wounds (Quirós and Nusrat 2018), were mostly represented in epithelial cells that do not respond to IFN-λ (**Figure 3A**).

**FIGURE 3.**
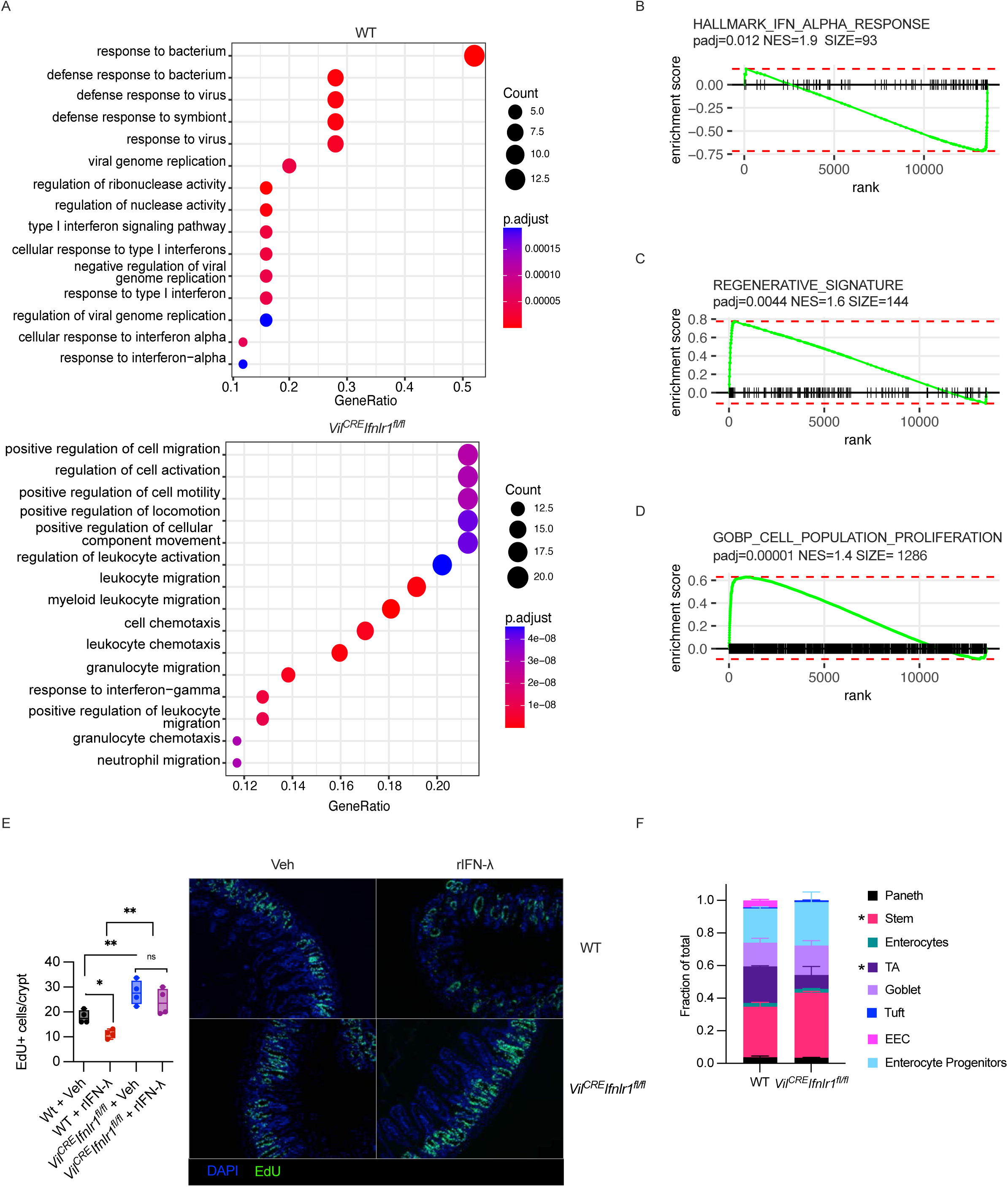
IFN-λ signaling induces an antiproliferative program in small intestine epithelia. (**A-D, F**) *Vil^CRE^ Ifnlr1*^fl/fl^ mice or WT mice received 11 Gy ionizing radiation, with lead shielding of the upper extremities. Targeted transcriptomics was performed on small intestinal crypts isolated 96h after irradiation. (**A**) Dot plots of Gene Ontology (GO) enrichment analysis. GO terms enriched in crypts from WT mice (*upper panel*) or *Vil^CRE^ Ifnlr1*^fl/fl^ mice (*lower panel*) are shown. Gene ratio (x axis), adjusted p-value (color) and gene count (dot size) are depicted. (**B-D**) GSEA enrichment plots of the HALLMARK_IFN_ALPHA_RESPONSE (**B**), REGENERATIVE SIGNATURE (as previously described (Yui, Azzolin et al. 2018)) (**C**), and the GO biological process Cell Population proliferation (“GOBP_CELL_POP_PROLIF) (**D**) are depicted. padj: adjusted p-value, NES: Normalized enrichment score, SIZE: size. (**E**) *Vil^CRE^ Ifnlr1*^fl/fl^ mice or WT mice were irradiated as in (**A-D**) and treated with either 50 mg kg^-1^day^-1^ of rIFN-λ (rIFN-λ), or saline vehicle (Veh). After 96h mice were pulsed with EdU for 2h. Number of EdU^+^ cells per crypt quantified by immunohistochemistry (IHC) (*left panel*), and representative IHC sections (*right panel*) are depicted. (**F**) Targeted transcriptomics data from WT or *Vil^CRE^ Ifnlr1*^fl/fl^ small intestinal crypts were deconvoluted based on publicly available single-cell RNA-seq (scRNA-seq) datasets (Haber, Biton et al. 2017) using CIBERSORTx (Newman, Steen et al. 2019) to extrapolate the relative cellular composition of samples. Paneth: Paneth cells; Stem: Intestinal stem cells; Enterocytes: small intestine enterocytes; TA: Transit amplifying cells; Goblet: Goblet cells; Tuft: tuft cells; EEC: Enteroendocrine cells; EP: Enterocyte progenitors. (**E**) Box plots are depicted. Each dot represents a mouse. Median, range and interquartile range are depicted. (**F**) Mean and SEM of 4 samples (WT) and 3 samples (*Vil^CRE^ Ifnlr1*^fl/fl^) are depicted. Statistics: (**E, F**) Two-way ANOVA with Šidak correction for multiple comparisons. ns= not significant (p > 0.05); *p < 0.05;**p < 0.01; ***p < 0.001; ****p < 0.0001.

Gene set enrichment analysis (GSEA) confirmed that genes associated with IFN responses were significantly enriched in WT, compared to knock-out, epithelial cells (**Figure 3B, S3A**). We next assessed the relative enrichment of a previously identified colitis-associated regenerative epithelial gene-set (Yui, Azzolin et al. 2018), as well as gene-sets associated with epithelial cell proliferation. Both gene-sets were significantly enriched when epithelial cells did not respond to IFN-λ (**Figure 3C, D, S3B, C**).

To assess whether IFN-λ-dependent delayed tissue restitution is characterized by reduced cell proliferation *in vivo*, we administered the thymidine analog 2’-deoxy-5-ethynyluridine (EdU) two hours before mice were euthanized and measured cell proliferation in either WT littermates or *Vil^CRE^Ifnlr1^fl/fl^* mice, administered or not rIFN-λ. We found that exogenous rIFN-λ reduced the number of EdU-positive cells per crypt in WT but not *Vil^CRE^Ifnlr1^fl/fl^* mice (**Figure 3E**). Also, that *Vil^CRE^Ifnlr1^fl/fl^*mice had a significant increased number of proliferating cells per crypts, compared to WT littermates, irrespectively of the administration of rIFN-λ (**Figure 3E**).

During tissue repair that follows radiation damage (Metcalfe, Kljavin et al. 2014) or colitis (VanDussen, Sonnek et al. 2019), specialized ISCs drive re-epithelialization by massively proliferating. Therefore, the decreased number of proliferating epithelial cells we observed in WT mice may reflect the lack of reparatory ISCs that proliferate. To assess whether endogenous IFN-λ affected the cellular composition of the small intestine in WT or *Vil^CRE^Ifnlr1^fl/fl^* mice that were irradiated, we used CIBERSORTx (Newman, Steen et al. 2019) and deconvoluted our bulk RNAseq data based on single-cell RNAseq data previously published (Haber, Biton et al. 2017). Our deconvolution analysis revealed that, while most epithelial cell types did not present major significant differences, the ISC compartment was significantly expanded in mice that were irradiated and whose epithelial cells do not respond to IFN-λ (**Figure 3F**). In keeping with retro- differentiation of transit-amplifying (TA) cells to replenish the ISC compartment upon ISC depletion (Wang, Chiang et al. 2019, Ohara, Colonna et al. 2022), TA cells were significantly decreased in the small intestine of *Vil^CRE^Ifnlr1^fl/fl^* mice, compared to WT controls (**Figure 3F**). The expansion of the Lrg5^+^ compartment in mice that do not respond to IFN-λ was confirmed by qPCR (**Figure S3D**). We also confirmed that the major epithelial cell populations analyzed were not different under homeostatic conditions in *Vil^CRE^Ifnlr1^fl/fl^*mice or WT littermates (**Figure S3E**). Overall, these data demonstrate that IFN-λ initiates a transcriptional program that reduces tissues restitution, limits ISC cell expansion, and, thus, dampens the overall capacity of epithelial cells to proliferate.

### IFN-λ controls the expression of ZBP1 and the activation of Gasdermin C

The reduced expansion of ISC can be driven either by increased cell death of ISCs and/or TA cells, reduced proliferative programs, or both. To determine the molecular mechanisms regulated by IFN-λ to dampen tissue restitution, we identified the genes that were significantly differentially regulated in epithelial cells derived from irradiated *Vil^CRE^Ifnlr1^fl/f^*^l^, compared to WT, mice (**Figure 4A**). As expected, multiple ISGs were among the genes significantly downregulated in cells that cannot respond to IFN-λ (**Figure 4A**). Intriguingly, *Zbp1* was among these genes. ZBP1 is a key component in the multiprotein complex PANoptosome, which encompasses effectors of several forms of cell death, and is an important regulator of cell fate (Kuriakose and Kanneganti 2018). We also found that protein levels of ZBP1, as well as another ISG such as RSAD2, were upregulated in epithelial cells of the small intestine upon *in vivo* administration of rIFN-λ in non-irradiated WT mice (**Figure S4A**). Upregulation of these proteins was prevented in epithelial cells derived from *Vil^CRE^Ifnlr1^fl/fl^*mice and was not different in the absence of rIFN-λ in the two backgrounds (**Figure S4A**). Among other genes significantly downregulated in cells that do not respond to IFN-λ there were two members of the gasdermin C (GSDMC) family. GSDMs are critical effectors of pyroptosis, a form of inflammatory cell death (Kovacs and Miao 2017). Compared to other GSDMs, very little is known about the functions of GSDMC, and scattered reports have involved GSDMC in the lytic death of tumor cells (Hou, Zhao et al. 2020, Zhang, Zhou et al. 2021), or of enterocytes during helminth infections (Xi, Montague et al. 2021). Of note, non-irradiated mice administered with rIFN-λ do not show upregulation of the GSDMC protein (**Figure S4A**), demonstrating that additional pathways associated with irradiation and/or tissue damage and repair regulate *Gsdmc* gene expression and/or protein synthesis.

**FIGURE 4.**
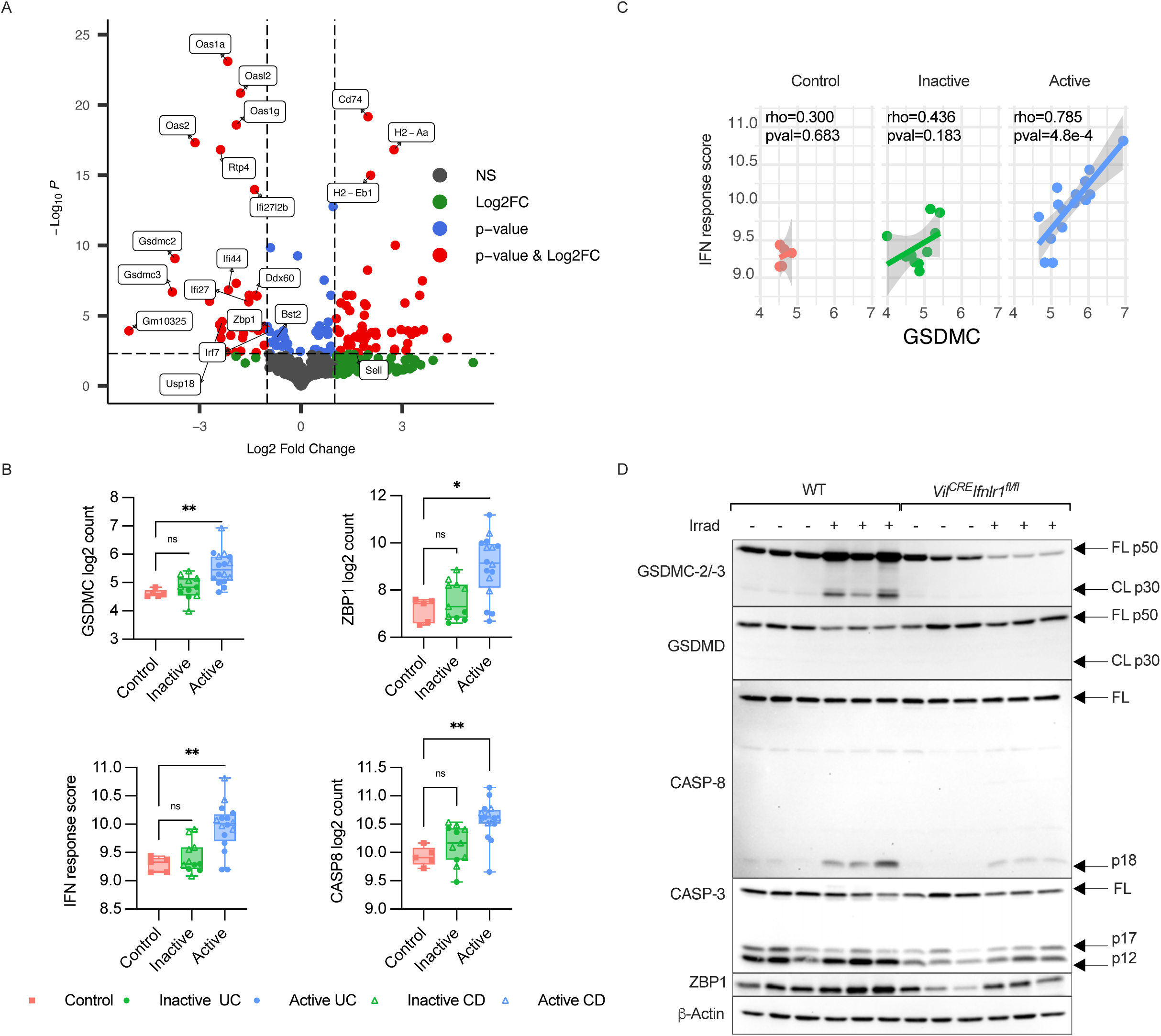
IFN-λ controls the expression of ZBP1 and the activation of gasdermin C. (**A**) *Vil^CRE^ Ifnlr1*^fl/fl^ mice or WT mice were irradiated as in Figure 2A. Targeted transcriptomics was performed on freshly isolated small intestinal crypts. Volcano plot depicting differentially expressed genes (DEGs) between *Vil^CRE^ Ifnlr1*^fl/fl^ and WT small intestinal crypts. DEGs (P< 0.005) with a fold change >2 (or <−2) are indicated in red; DEGs with a fold change <2 (or >−2) are indicated in blue. Nonsignificant DEGs (P> 0.005) and genes not differentially expressed are indicated in green and gray, respectively. Positive values represent genes overexpressed in *Vil^CRE^ Ifnlr1*^fl/fl^, negative values represent genes overexpressed in WT. (**B**) RNA sequencing was performed on colon biopsies from control patients, IBD patients with inactive disease, and IBD patients with active disease, (see Materials and Methods). Square symbols represent controls, round symbols represent ulcerative colitis (UC) patients, triangles represent Crohn’s disease patients (CD). Box plots with median, range and interquartile range are depicted. Each symbol represents one patient. Expression of *GSDMC* (*top left panel*), *ZBP1* (*top right panel*), *CASP8* (*bottom right panel*) expressed as normalized log2 count is depicted. Mean expression of genes belonging to the GSEA HALLMARK_IFN _ALPHA_RESPONSE gene set (IFN Response score), is depicted (*bottom left panel*). (**C**) Dot plot depicting the correlation between GSDMC expression and the IFN Response score performed on the same samples as (**B**). Each point represents a patient, solid lines represent linear regression, shaded area depicts the confidence interval. Spearman correlation coefficient (rho) and the relative p-value (pval) are indicated for each graph. (**D**) Small intestinal crypts were isolated from *Vil^CRE^ Ifnlr1*^fl/fl^ mice or WT mice irradiated as in Figure 2A (Irrad +) or not (Irrad -). Immunoblot analysis of the indicated proteins was performed. GSDMC-2/-3 FL p50: full length 50 kDa GSDMC-2/3; GSDMC-2/-3 CL p30: N-terminal 30 kDa cleaved protein; GSDMD FL p50: full length GSDMD 50 kDa; GSDMD CL p30: N-terminal 30 kDa cleaved protein; CASP-8 FL: full length CASP-8; CASP-8 p18: 18 kDa CASP-8 cleavage fragment; CASP-3 FL: full length CASP-3; CASP-3 p17: 17 kDa CASP-3 cleavage fragment; CASP-3 p12: 12 kDa CASP-3 cleavage fragment. Each lane represents one mouse. Representative data of 3 independent experiments is depicted. Statistics: (**B**) Kruskal Wallis test with Dunn correction for multiple comparisons was performed. ns= not significant (p > 0.05); *p < 0.05;**p < 0.01; ***p < 0.001; ****p < 0.0001.

IBD patients present increased levels of IFN-λ and/or *Ifnlr1* (Chiriac, Buchen et al. 2017, Gunther, Ruder et al. 2019). Prompted by our findings in irradiated mice, we assessed the expression levels of *ZBP1* and *GSDMC* (the only GSDMC present in humans) in the biopsy derived from a cohort of IBD patients with active or inactive disease, or non-IBD controls (see Material and Methods for details). We found a significant increase in the expression of *ZBP1* as well as *GSDMC* in patients with active IBD, compared to non-IBD controls (**Figure 4B**). Similar expression trends were observed in RNAseq datasets derived from rectal mucosal biopsies from ulcerative colitis (UC) pediatric patients (PROTECT cohort) and from ileal biopsies from Crohn’s disease (CD) pediatric patients (RISK cohort) in two independent cohorts previously published (Haberman, Tickle et al. 2014, Haberman, Karns et al. 2019) (**Figure S4B**). In keeping with a key role of IFNs also in IBD patients, we found that genes regulated in response of IFNs (evaluated as mean expression of genes that belong to the “HALLMARK _IFN_ALPHA_RESPONSE” gene- set (Liberzon, Birger et al. 2015) and indicated as “IFN response score”) were significantly enriched in IBD patients with active disease, compared to controls or patients with inactive disease (**Figure 4B**). In support of a role of IFN-λ in modulating GSDMC expression, *GSDMC* levels positively correlated with the IFN response score in patients with active disease, but not in the other subjects analyzed (**Figure 4C**). These data indicate that IFN induction and upregulation of the genes that encode for ZBP1 and GSDMC are hallmarks of intestinal damage both in mice and humans.

GSDMs exert their pyroptotic function upon cleavage by caspases (Kovacs and Miao 2017), when the N-terminal cleavage product oligomerizes to form lytic pores in the cell membrane, leading to the loss of ionic homeostasis and cell death (Broz, Pelegrín et al. 2020). We, thus, tested whether GSDMC-2/-3 were cleaved in epithelial cells of the small intestine upon irradiation and confirmed that irradiated, but not non-irradiated, mice not only showed increased levels of GSDMC-2/-3, but also that GSDMC-2/-3 were efficiently cleaved in WT, but not *Vil^CRE^Ifnlr1^fl/fl^*, mice (**Figure 4D**). In contrast, another key effector of pyroptosis, GSDMD, was not activated.

GSDMC is primarily cleaved by Caspase-8 (Hou, Zhao et al. 2020, Zhang, Zhou et al. 2021). Indeed, the pattern of bands of cleaved GSDMC-2/-3 is compatible with the activity of Caspase-8 (Julien and Wells 2017). We thus investigated the activation of Caspase-8. Caspase- 8 was not activated in WT or *Vil^CRE^Ifnlr1^fl/fl^* mice in the absence of irradiation (**Figure S4A**). In contrast, epithelial cells derived from irradiated WT littermates, but not *Vil^CRE^Ifnlr1^fl/fl^* mice, efficiently activated Caspase-8 (**Figure 4D**). Of note, we found that expression of *CASP8* was increased in patients with active IBD compared to non-IBD controls (**Figure 4B, S4B**). In keeping with the capacity of ZBP1 to control the activation of multiple caspases (Kuriakose and Kanneganti 2018), we found that a similar pattern of activation was also true for Caspase-3 (**Figure 4D**).

Overall, these data demonstrate that IFN-λ initiates in the small intestine of irradiated mice a signaling cascade that allows the upregulation of ZBP1, the activation of Caspase-8/-3, and the induction and cleavage of GSDMC, an executor of pyroptosis. Also, that similar programs are transcriptionally upregulated in IBD patients with active disease.

### The ZBP1-GSDMC axis induced by IFN-λ controls epithelial cell death and proliferation

To assess directly the role of the signaling cascade initiated by IFN-λ in driving cell death, we used intestinal organoids. Mouse and human small intestinal organoids seeded in the presence of rIFN-λ died between 48 and 72h from treatment (**Figure 5A, S5A**). Dying cells assumed typical changes associated with pyroptosis including swelling and sudden disruption of the plasma membrane and liberation of nuclear DNA (**Figure 5A**). By using organoids derived from WT or *Stat1*^-/-^ mice, we also confirmed that gene transcription induced by IFN-λ was necessary to induce cell death (**Figure 5B**). No differences were observed between the two genotypes in the absence of rIFN-λ (**Figure 5B**). Similar to what we observed in irradiated epithelial cells *in vivo*, we found that rIFN-λ administration to small intestine organoids profoundly diminished the level of *Lgr5* expression, suggesting a defect in the maintenance and/or proliferation of ISCs (**Figure S5B**). In agreement with a reduced number of ISCs that proliferate, we also found that cell proliferation (as measured by EdU incorporation) was significantly decreased in IFN-λ-treated mouse, as well as human organoids (**Figure 5C, S5C**).

**FIGURE 5.**
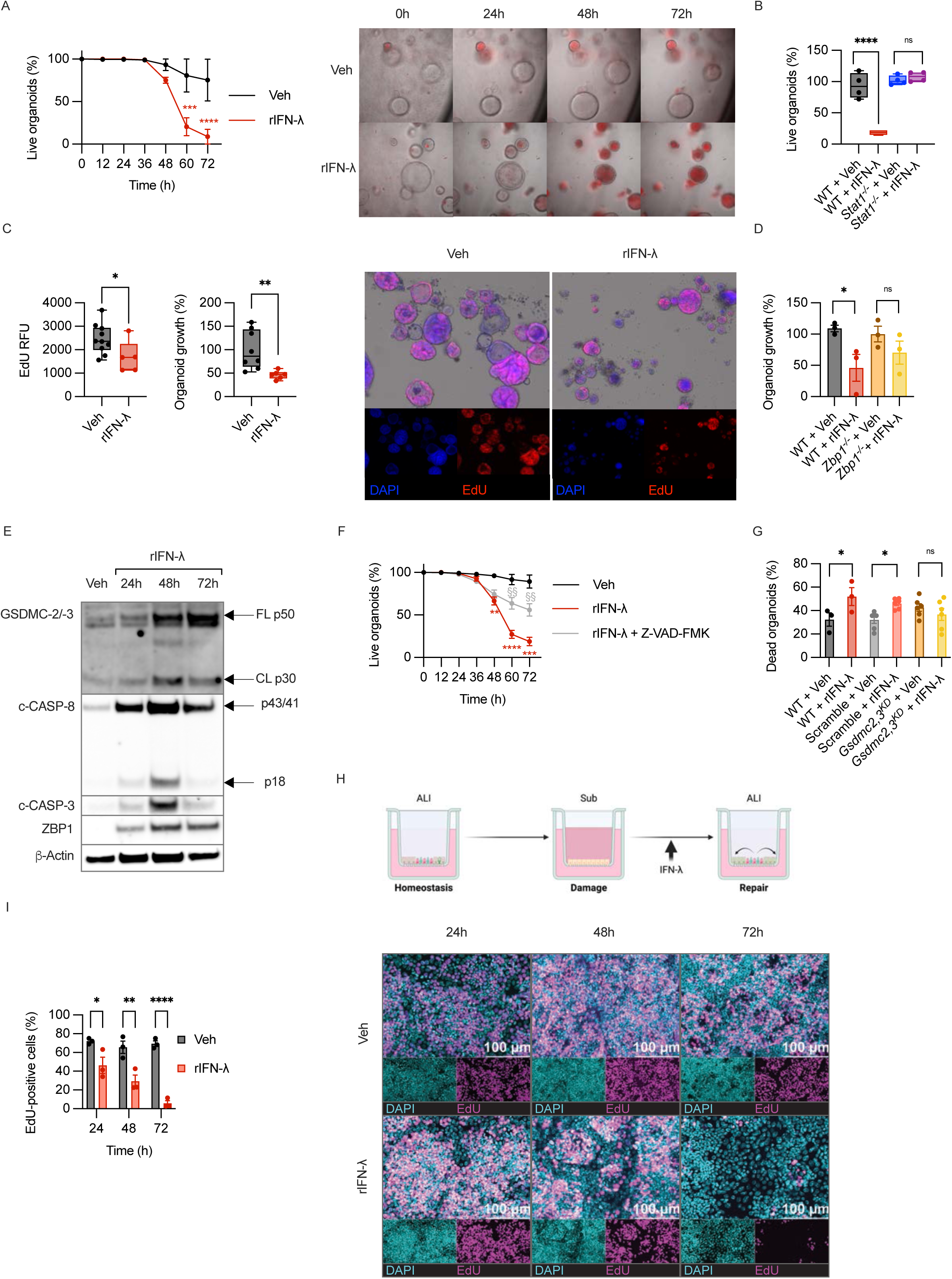
IFN-λ inhibits epithelial proliferation and survival in intestinal organoids *in vitro*. (**A**) Mouse small intestinal organoids were seeded from freshly isolated crypts and allowed to grow for 48h. Organoids were then treated with 200ng/ml of rIFN-λ in the presence of 1ug/ml propidium iodide (PI) and imaged every 12h over 72h. Percentage of live organoids (*left panel*) was calculated as percentage of PI^-^ organoids over the total number of live organoids in each well. Representative image of 3 independent experiments is depicted (*right panel*). (**B**) Small intestinal organoids derived from WT or *Stat1^-/-^* mice were seeded and treated as in (**A**). Percentage of live organoids was calculated as percentage of PI^-^ organoids over the total number of live organoids in each well. (**C**) Small intestinal organoids were seeded and treated as in (**A**) for 72h. Organoids were pulsed with EdU for 6h. Organoids were stained for EdU incorporation (RED), and DAPI (BLUE). Mean fluorescence of EdU staining (*left panel*), relative organoid growth (*middle panel*) and representative images (*right panel*), are depicted. The relative growth of organoids is measured as the % of their area over untreated control organoids. (**D**) Small intestinal organoids derived from WT or *Zbp1^-/-^* mice were seeded and treated with 200ng/ml of rIFN-λ and imaged at 48h. The relative growth of organoids measured as in (**C**) is depicted. (**E**) Small intestinal organoids were treated as in (**A**) for 24, 48 and 72h. Immunoblot analysis of the indicated proteins was performed. GSDMC-2/-3 FL p50: full length 50kDa GSDMC-2/-3; GSDMC- 2/-3 CL p30: N-terminal 30kDa cleaved protein; c-CASP-8 p43: 43 kDa CASP-8 cleavage fragment; c-CASP-8 p18: 18 kDa CASP-8 cleavage fragment. Representative blot of three independent experiments. (**F**) Small intestinal organoids were seeded as in (**A**) and either treated with 200ng/ml of rIFN-λ alone (rIFN-λ) or with rIFN-λ in the presence of the pan caspase inhibitor Z-VAD-FMK (40uM). Organoids were then followed for 72h and % of live organoids was evaluated as in (**A**). Statistical comparison between “rIFN-λ” and “Veh” are depicted in red (*), comparison between “rIFN-λ + Z-VAD-FMK” and “rIFN-λ” is depicted in grey (§). (**G**) Small intestinal organoids were either left untreated (WT) or infected with a lentivirus expressing GFP and either a Gasdermin-2/Gasdermin-3 targeting (*Gsdmc2, 3^KD^*) small harpin (sh)RNA, or a scrambled control shRNA (Scramble). Organoids were grown for 5 days and then treated with 200ng/ml of rIFN-λ or vehicle control for 48h. Survival of WT controls or lentiviral-infected GFP^+^ cells was assessed by cytofluorimetry by staining with Zombie dye and calcein. % of dead cells represent cells positive for Zombie dye staining and negative for calcein. (**H**) Experimental scheme for the establishment of 2D Air Liquid Interface (ALI) organoid cultures and modeling of damage and repair responses. Organoids were seeded in transwells and grown to confluence. The apical side was then exposed to air up to 14 days, which favored differentiation of a homeostatic monolayer. Organoids were then submerged for 7 days to induce damage responses. After 7 days they were re-exposed to air to stimulate repair responses. Concomitantly with re-exposure to air, organoids were treated with 200ng/ml of rIFN-λ for 3 days. (**I**) Organoids treated as in (**H**) were pulsed with EdU for 2h to mark proliferating cells. Quantification of the percentage of EdU^+^ cells per field of view (*lelft panel*) and representative images (*right panel*) are depicted. Edu (MAGENTA), and DNA stain DAPI (CYAN) are depicted. Representative image of 3 independent experiments. (**A, F**) Mean and SEM of 3 (**A**) and 5 (**F**) biological replicates per group are depicted. (**B, C**) Box plots are depicted. Each dot represents a biological replicate. Median, range and interquartile range are depicted. (**D, G, I**) Scatter plots are depicted. Each dot represents a biological replicate. Statistics: (**A, F**) Two-way ANOVA with Tukey correction for multiple comparisons. (**B, C, D, G, I**) Two-way ANOVA with Šidak correction for multiple comparison. (**C**) Unpaired t test. ns= not significant (p > 0.05); *or §p < 0.05;**or §§p < 0.01; ***or §§§p < 0.001; ****or §§§§p < 0.0001.

To assess the involvement of ZBP1 in these processes, we derived organoids from either WT or *Zbp1*^-/-^ mice. Organoids were differentiated in the presence or absence of IFN-λ and their survival and/or growth was followed for 72h. Survival and growth of organoids, derived either from the small or large intestine, differentiated from WT, but not *Zbp1*^-/-^, mice were significantly reduced upon the administration of rIFN-λ (**Figure 5D, S5D**). In agreement with the capacity of IFN-λ to activate a ZBP1/Caspase-8/GSDMC axis, organoids grown for 6 days and then administered with rIFN-λ showed ZBP1 upregulation and cleavage of GSDMC-2/-3 and Caspase-8 and Caspase-3 (**Figure 5E**). Furthermore, inhibition of caspase activity with the pan-caspase inhibitor Z-VAD-FMK protected the organoids from cell death induced by IFN-λ (**Figure 5F**). To directly assess the involvement of GSDMC in this process, we knocked down *Gsdmc-2 and Gsdmc-3* in small intestine organoids and found that, similar to *Zbp1*^-/-^ organoids, upon exposure to rIFN-λ, survival of organoids that do not express *Gsdmc2, 3* was significantly increased compared to controls (**Figure 5G**).

To better reflect the cycles of injury and repair characteristic of IBD and mouse models of colitis, we implemented a previously described model of long-term organoid culture (Wang, Chiang et al. 2019). We grew organoids in a two-dimensional (2D) epithelial monolayer system and exposed their apical side to air, to obtain a self-organizing monolayer that mimics cells in homeostasis. This monolayer can then be re-submerged in medium (to elicit damage response mimicking *in vivo* epithelial injuries) and re-exposed to air (which induces epithelial regeneration responses) (**Figure 5H**). When we treated with IFN-λ the epithelial monolayer after re-exposure to air, the proliferative repair response was curbed, as demonstrated by the failure to incorporate EdU (**Figure 5I**).

Overall, our data demonstrate that IFN-λ signaling induces a ZBP-1-GSDMC axis that controls epithelial cell survival, and dampens the capacity of ISCs to proliferate and orchestrate tissue restitution.

## Discussion

In our work, we revealed that IFN-λ restrains the restitution of the intestinal mucosae secondary to either inflammatory damage or ionizing radiations toxicity. We reveal the capacity of IFN-λ to initiate a previously overlooked molecular cascade in intestinal epithelial cells that allows the induction of ZBP1, the activation of caspases, and the induction and cleavage of GSDMCs. We also found that similar pathways are transcriptionally upregulated in IBD patients with active disease. Induction of epithelial cell death via the ZBP1-GSDMC axis reduces the number of ISCs, and dampens the proliferation and restitution of epithelial cells, thus affecting the re-epithelization of the injured intestine. Finally, we revealed that IFN-λ, but not type I IFNs, are the major drivers of the delayed restitution *in vivo* in mouse models of gut damage.

The immune system is endowed with the capacity not only to protect against pathogen invasion but also to maintain tissue homeostasis. Fundamental to exert these activities, is the fine balance between anti-microbial functions of the immune system that can drive tissue damage, and the regenerative capacity of organs and tissues. Many cellular and molecular mediators of the immune system are involved in exerting anti-microbial and potentially damaging functions, but several can also modulate mucosal repair. Here, we focused our attention on a group of IFNs, known as type III IFNs or IFN-λ. IFN-λ activities at mucosal surfaces are essential to limit pathogen spread while reducing inflammation and immune cell infiltration (Broggi, Granucci et al. 2020). IFN-λ and type I IFNs regulate very similar transcriptional programs, but the limited expression of the IFNLR restricts the activity of IFN-λ to epithelial cells, hepatocytes, neutrophils and few other cell types and, thus, reduces the extent of the inflammation (Broggi, Granucci et al. 2020). The limited number of cells that respond to IFN-λ signaling, and the reduced capacity of IFN-λ, compared to type I IFNs, to activate IRF1 (Forero, Ozarkar et al. 2019) allow to preserve the functionality of mucosal tissues during an immune response. Although the protective functions of IFN-λ in the gut, and in general at mucosal surfaces, are well known (Broggi, Granucci et al. 2020), much less is known about the functions of this group of IFNs during the healing phase that follows intestinal tissue damage. Our data reveal the unique capacity of IFN-λ, compared to type I IFNs, to negatively affect tissue restitution in the intestine, possibly opening new ways of therapeutic intervention for individuals that encounter tissue damage such as IBD patients or subjects exposed to radiation therapies.

A common feature revealed by our analyses is that IFN-λ affects the survival of the cells, decreases the number of ISCs, and dampens the proliferation of the cells in the crypts, both *in vivo* in mice and *in vitro* in both human and mouse organoids. Tissue restitution in the gut is regulated by a complex crosstalk between epithelial cells, immune cells, microbial stimuli, and mesenchymal cells and culminates in the proliferation of ISCs. Treatment with ionizing radiation induces widespread epithelial damage and targets in particular proliferating ISCs in the intestinal crypt, making it an ideal model to understand the dynamics of ISCs proliferation and intestinal healing. Lrg5^+^ ISCs support normal cell turnover as well as injury-induced restitution (Metcalfe, Kljavin et al. 2014). When ISC are depleted by radiation, or by immune-mediated tissue-damaging events, TA cells retro-differentiate and acquire new stem-like properties in the small, as well as in the large, intestine (Wang, Chiang et al. 2019, Ohara, Colonna et al. 2022). These cells then proliferate to allow the re-epithelization of the damaged tissue. We and others previously described the capacity of IFN-λ to dampen lung epithelial cell proliferation (Broggi, Ghosh et al. 2020, Major, Crotta et al. 2020) and to instruct anti-proliferative transcriptional programs in the lung of patients infected with SARS-CoV-2 (Sposito, Broggi et al. 2021). Nevertheless, our new findings in the gut suggest that decreased proliferation assessed at transcriptional and cellular levels is due to augmented cell death, possibly occurring in newly generated ISCs or in TA cells. If similar processes also take place in the lung, it remains an open question that will require further investigation.

Another interesting observation we made is that exogenous administration of IFN-λ does not induce caspase or GSDMC activation *in vivo* in the absence of tissue damage, although it induces ZBP1 upregulation at the transcriptional as well as protein level. ZBP1 is a Z-DNA binding protein, and is part of the PANoptosome, a multiprotein complex that governs the cell fate (Place, Lee et al. 2021). PANoptosis is a form of cell death that encompasses pyroptosis, apoptosis, and necroptosis. ZBP1 can interact directly or indirectly with proteins that regulate cell death and drive the activation of apoptotic caspases 8, 3 and 7, the necroptosis effector MLKL, or pyroptosis effectors Casp-1, 11 and GSDMD. So far, GSDMC was not associated with ZBP1 and/or PANoptosis. Our data highlight the existence of a ZBP1-GSDMC axis that appears to be the preferential pathway of cell death that is active during cycles of intestinal epithelial damage and restitution. Upregulation of ZBP1 alone is not sufficient to trigger the full activation of the PANoptosome, which is consistent with our inability to detect toxic effects of IFN-λ in the absence of inflammation or tissue damage. Conversely, ZBP1 can be activated both by binding microbial-derived nucleic acids (Kuriakose, Zheng et al. 2018, Muendlein, Connolly et al. 2021) or by binding host-derived Z-DNA following oxidative damage of the mitochondria (Szczesny, Marcatti et al. 2018). It is, thus, possible that *in vivo*, under tissue-damaging conditions, either microbiota- or host-derived DNA becomes available to induce the assembly and activation of the PANoptosome downstream of ZBP1. In contrast to our *in vivo* data, administration of IFN-λ to murine or human intestinal organoids induces the ZBP1-GSDMC axis and drives cell death. We, thus, speculate that under our *in vitro* experimental conditions a “tissue damage” signal, e.g. Z-DNA from dying cells that are differentiating *in vitro*, is available, thus making additional signals unnecessary.

Linked to the above-mentioned observations, we also found that the ISC compartment is not altered under homeostatic conditions in *Vil^CRE^Ifnlr1^fl/fl^* mice compared to WT mice, suggesting that the basal level of IFNs present during homeostasis (Van Winkle, Peterson et al. 2022) does not affect the normal turnover of gut epithelial cells. Intriguingly, it has been recently shown that IFN-λ-dependent responses at homeostasis are restricted to pockets of mature enterocytes in the small intestine and in the colon (Van Winkle, Peterson et al. 2022). In contrast, when mice are injected with rIFN-λ, or infected with murine rotavirus, a type of virus potently controlled by IFN-λ (Walker, Sridhar et al. 2021), responses to IFN-λ broadly distribute along the epithelial layer. These data, together with our new findings, suggest that the compartmentalization of IFN-λ signaling at homeostasis preserves the functions of ISCs and the normal turnover of gut epithelial cells.

Our models of intestinal damage either of the colon, in the DSS-colitis model, or of the small intestine, in the radiation model, highlight the centrality of IFN-λ and its capacity to delay tissue restitution. Two previous studies suggested that IFN-λ may play an opposite role and favor tissue restitution during colitis (Chiriac, Buchen et al. 2017, McElrath, Espinosa et al. 2021). Nevertheless, both studies were performed by inducing colitis in total *Ifnlr1*^-/-^ mice and thus they suffer the confounding activity of IFN-λ on neutrophils. The absence of IFN-λ signaling in neutrophils potentiates tissue damage (Broggi, Tan et al. 2017), making it hard to compare the tissue restitution phase with WT mice that start from a different level of damage. Indeed, we always administered or blocked IFNs in our colitis model after the peak of the inflammatory phase. Alternatively, we used mice deficient for the IFNLR only in epithelial cells. Total knock-out mice were solely used in the radiation model in which damage and/or inflammation are not driven by neutrophils but by the ionizing radiations. The compartmentalized activity of IFN-λ in different cell types appears to be, thus, a key feature of this group of IFNs.

Overall, our data unveiled a new axis between IFN-λ, ZBP1 and GSDMC that governs tissue restitution in the gut and open new perspectives to future therapeutic interventions.

## Acknowledgements

IZ is supported by NIH grants 1R01AI121066, 1R01DK115217, 1R01AI165505 and contract no. 75N93019C00044, Lloyd J. Old STAR Program CRI3888, and holds an Investigators in the Pathogenesis of Infectious Disease Award from the Burroughs Wellcome Fund. AB is supported by the excellence initiative of Aix Marseille Université-A*Midex, a French “investissements d’Avenir” program: AMX-20-CE-01; the FRM amorçage de jeunes equipes grant AJE202010012468; and the ANR-JCJC grant “INTERMICI” ANR-21-CE15-0022. The French National Research Agency through the “Investments for the Future” program (France-BioImaging, ANR-10-INBS-04) contributed to support this work. We thank the imaging core facility (ImagImm) of the Centre d’Immunologie de Marseille-Luminy (CIML).

## Author contributions

BS designed, performed, analyzed the experiments; JM performed and analyzed murine organoid experiments; KBG and JT participated to experiments with human organoids and performed and analyzed the knock-down experiment; LS performed the RNAseq data analyses and the transcriptome analysis of IBD patients; NA participated to the histological analyses and *to in vitro* experiments; FG and PN performed the studies on IBD patients; VM and SBS gave advice on *in vitro* and *in vivo* experiments and analyzed human data from publicly available databases; AB conceived the project, designed and performed the experiments, supervised the study, and edited the paper; IZ conceived the project, designed the experiments, supervised the study, and wrote the paper. All authors reviewed and provided input to the manuscript. All authors declare no conflicts of interest.

## SUPPLEMENTARY FIGURE LEGENDS

**FIGURE S1.**
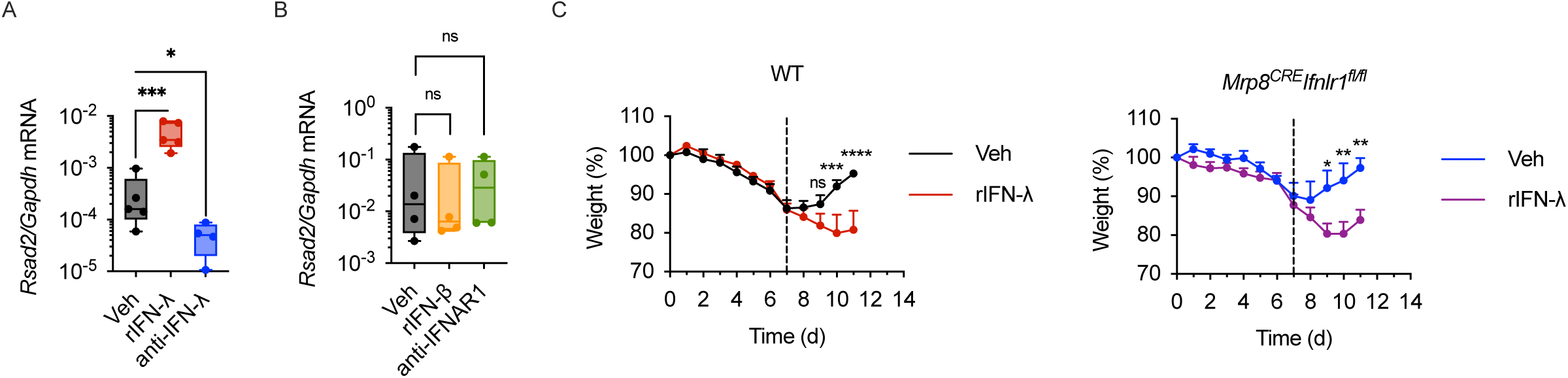
IFN-λ inhibits tissue recovery after DSS-colitis. **(A)** WT mice were treated with 2.5% DSS for 7 days. Upon DSS withdrawal mice were injected i.p. with either 50 µg kg-1day-1 of rIFN-λ or 12.5 mg kg^-1^day^-1^ of anti-IFN-λ2,3 antibody for five days. *Rsad2* relative mRNA expression in colonocytes on Day 14 is depicted. (**B**) Mice were treated with DSS as in (**A**). Upon DSS withdrawal mice were injected i.p. with either 50 mg kg^-1^day^-1^ of rIFN-β or 12.5 mg kg^-1^day^-1^ of anti-IFNAR1 antibody. *Rsad2* relative mRNA expression in colonocytes on Day 14 is depicted. (**C**) WT (*left panel*) or *Mrp8^CRE^Ifnlr1*^fl/fl^ mice (*right panel*) were treated with 2.5% DSS for seven days. Upon DSS withdrawal mice were injected i.p. with 50 µg kg^-1^day^-1^ of rIFN-λ. Weight is depicted. (**A, B**) Box plots are depicted. Each dot represents a mouse. Median, range and interquartile range are shown. (**C**) Mean and SEM of 5 mice per group are depicted. Statistics: (**A, B**) One-way ANOVA with Dunnett correction for multiple comparisons. (**C**) Two-way ANOVA with Tukey correction for multiple comparisons. ns= not significant (p > 0.05); *p < 0.05;**p < 0.01; ***p < 0.001; ****p < 0.0001.

**FIGURE S2.**
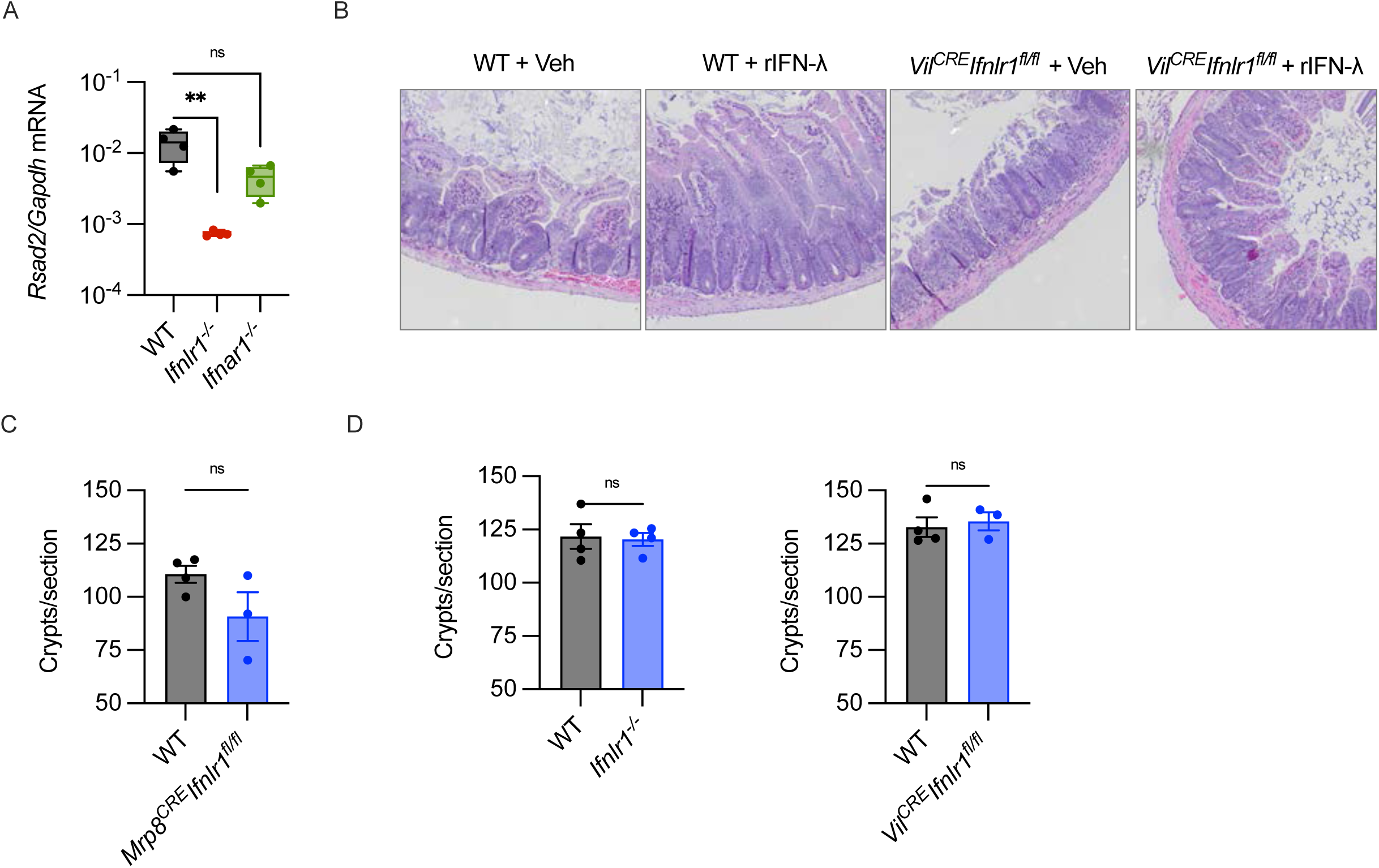
IFN-λ impairs epithelial regeneration after radiation damage. (**A**) WT, *Ifnlr1^-/-^*, or *Ifnar1^-/-^*mice received 11 Gy ionizing radiation, with lead shielding of the upper body. *Rsad2* relative mRNA expression in small intestinal crypt cells was evaluated 96h after irradiation. (**B**) *Vil^CRE^ Ifnlr1*^fl/fl^ mice or WT mice were irradiated as in (**A**) and treated with either 50 mg kg^-1^day^-1^ of rIFN-λ (rIFN-λ), or saline vehicle (Veh). Tissue repair in the small intestine was evaluated 96h after irradiation. Representative histological images of the small intestine are depicted. (**C**) WT mice and *Mrp8^CRE^Ifnlr1^fl/fl^*were irradiated as in (**A**). Tissue repair in the small intestine was evaluated 96h after irradiation by counting the number of intact crypts per histological section (Crypts/section). (**D**) Number of small intestinal intact crypts per histological section (Crypts/section) was evaluated in WT and *Ifnlr1^-/-^* (*left panel*) and WT and *Vil^CRE^Ifnlr1^fl/fl^* (*right panel*) mice at homeostasis. (**A**) Box plots are depicted. Each dot represents a mouse. Median, range and interquartile range are depicted. (**C, D**) Scatter plots are depicted. Each dot represents a mouse. Mean with SEM are depicted. Statistics: (**A**) One-way ANOVA with Dunnett correction for multiple comparisons. (**B**) Unpaired t test. ns= not significant (p > 0.05); *p < 0.05;**p < 0.01; ***p < 0.001; ****p < 0.0001.

**FIGURE S3.**
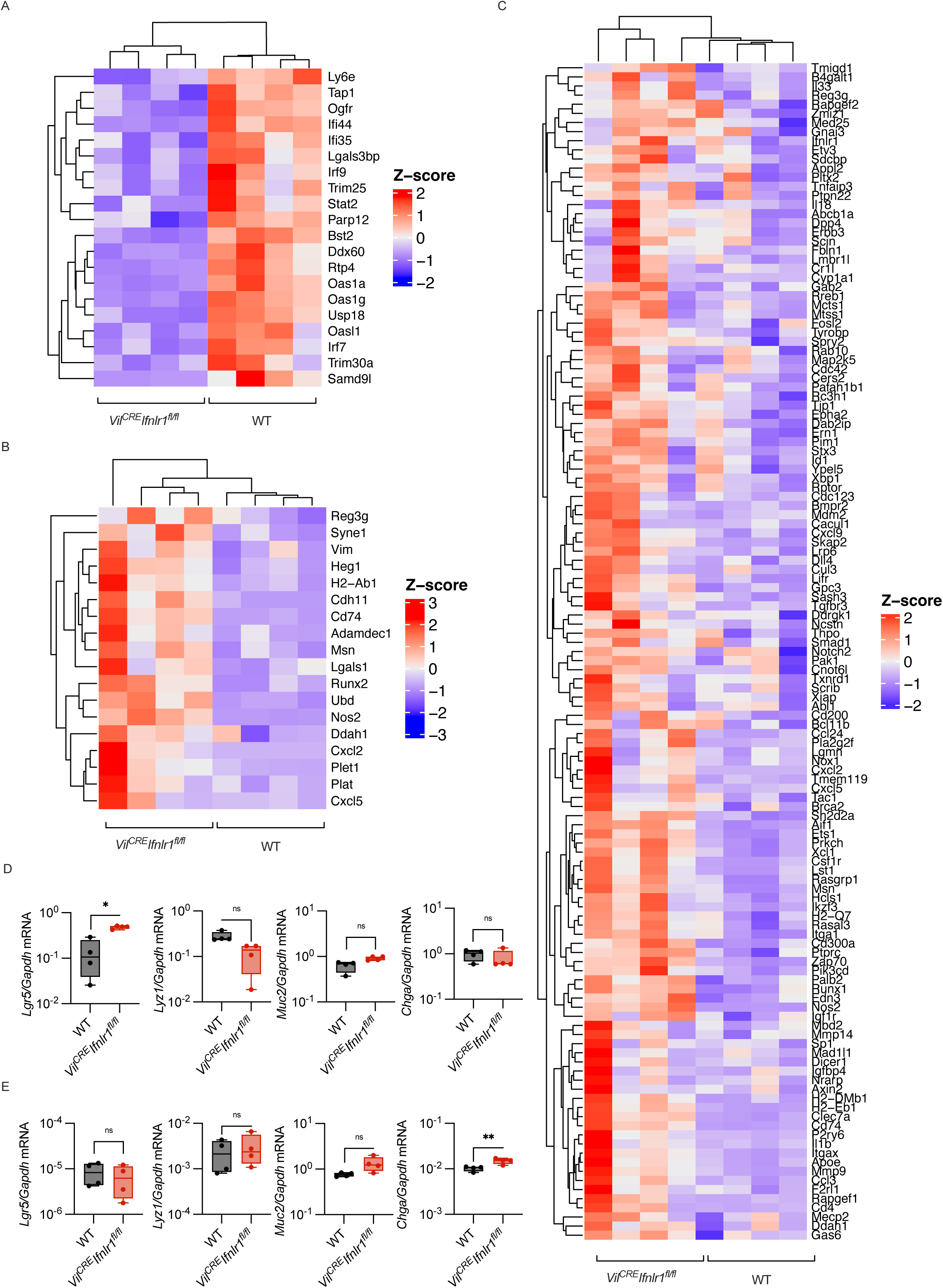
IFN-λ signaling induces an antiproliferative program in small intestine epithelia. (**A-D**) *Vil^CRE^ Ifnlr1*^fl/fl^ mice or WT mice received 11 Gy ionizing radiation, with lead shielding of the upper body. (**A, B**) Targeted transcriptomics was performed on small intestinal crypts isolated 96h after irradiation. (**A**) Heatmap depicting expression of genes in the leading edge of the enriched HALLMARK_IFN_ALPHA_RESPONSE gene set. The color is proportional to the Z Score. (**B**) Heatmap depicting expression of genes in the leading edge of the REGENERATION SIGNATURE gene set. The color is proportional to the Z Score. (**C**) Heatmap depicting expression of genes in the leading edge of the enriched GOBP_CELL_POPULATION_PROLIFERATION gene set. The color is proportional to the Z Score. (**D**) *Lgr5, Lyz1, Muc2, Chga* relative mRNA expression in small intestinal crypt cells was evaluated 96h after irradiation. (**E**) *Lgr5, Lyz1, Muc2, Chga* relative mRNA expression in small intestinal crypt cells isolated from *Vil^CRE^ Ifnlr1*^fl/fl^ mice or WT mice at homeostasis was evaluated. (**D, E**) Box plots are depicted. Each dot represents a mouse. Median, range and interquartile range are depicted. Statistics: (**D, E**) Unpaired t test. ns= not significant (p > 0.05); *p < 0.05;**p < 0.01; ***p < 0.001; ****p < 0.0001.

**FIGURE S4.**
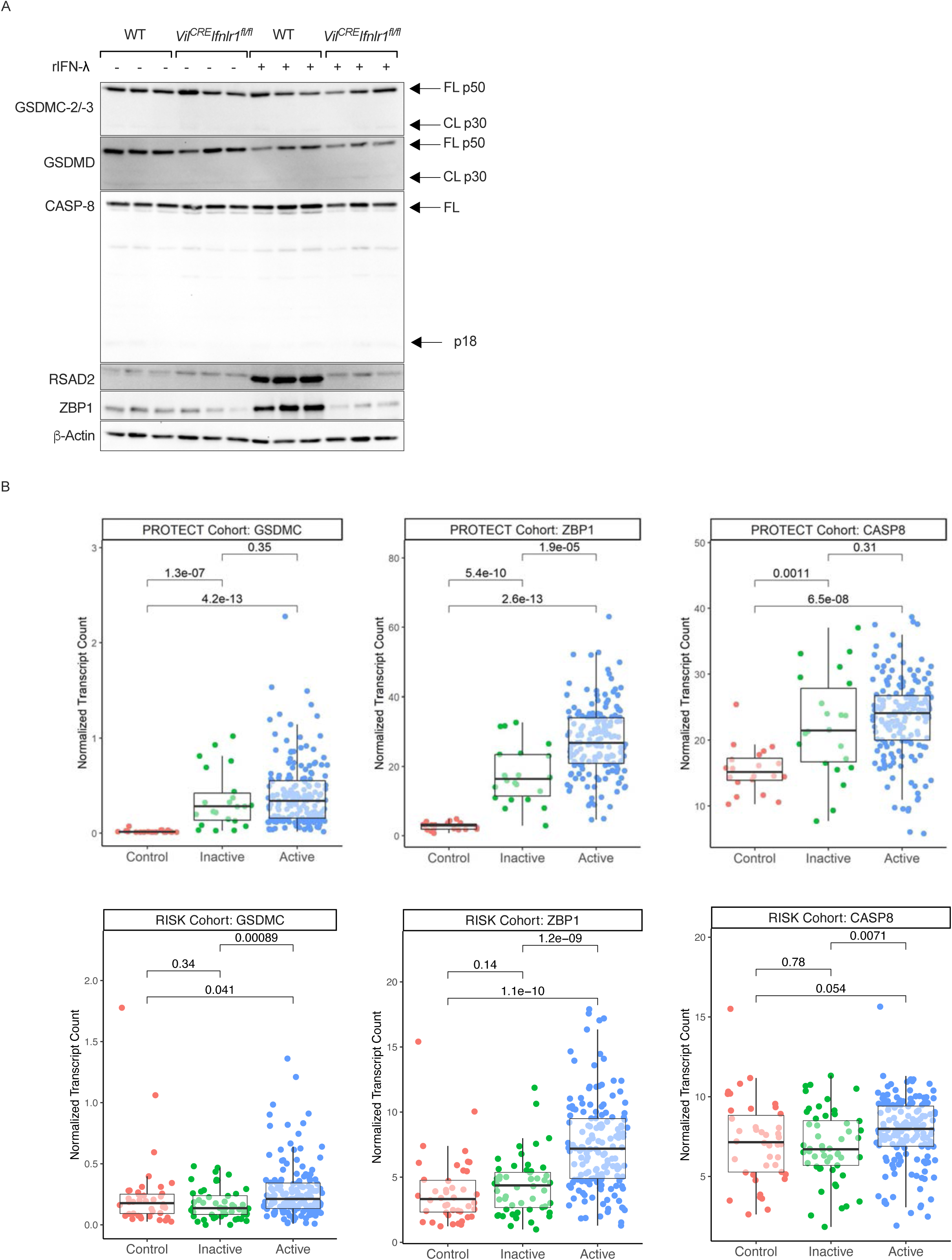
IFN-λ controls the expression of ZBP1 and the activation of Gasdermin C. (**A**) *Vil^CRE^ Ifnlr1*^fl/fl^ mice or WT mice were treated with either 50 mg kg^-1^day^-1^ of rIFN-λ (rIFN-λ), or saline vehicle (Veh). Immunoblot analysis of the indicated proteins was performed. Each lane represents one mouse. GSDMC-2/-3 FL p50: full length 50 kDa GSDMC-2/-3; GSDMC-2/-3 CL p30: N-terminal 30 kDa cleaved protein; GSDMD FL p50: full length 50 kDa GSDMD; GSDMD CL p30: N-terminal 30 kDa cleaved protein; CASP-8 FL: full length CASP-8; CASP-8 p18: 18 kDa cleavage fragment. (**B**) Expression (defined by normalized transcript counts; Transcripts Per Kilobase Million [TPM] for PROTECT cohort and Reads Per Kilobase Million [RPKM] for RISK cohort) of the indicated genes was assessed in bulk RNA-seq data from the PROTECT (pediatric UC) and RISK (pediatric ileal CD) cohorts and comparisons made between control patients, uninflamed IBD and inflamed IBD patients. P-values are based on non-parametric t-testing between assessed groups (Wilcoxon test).

**FIGURE S5.**
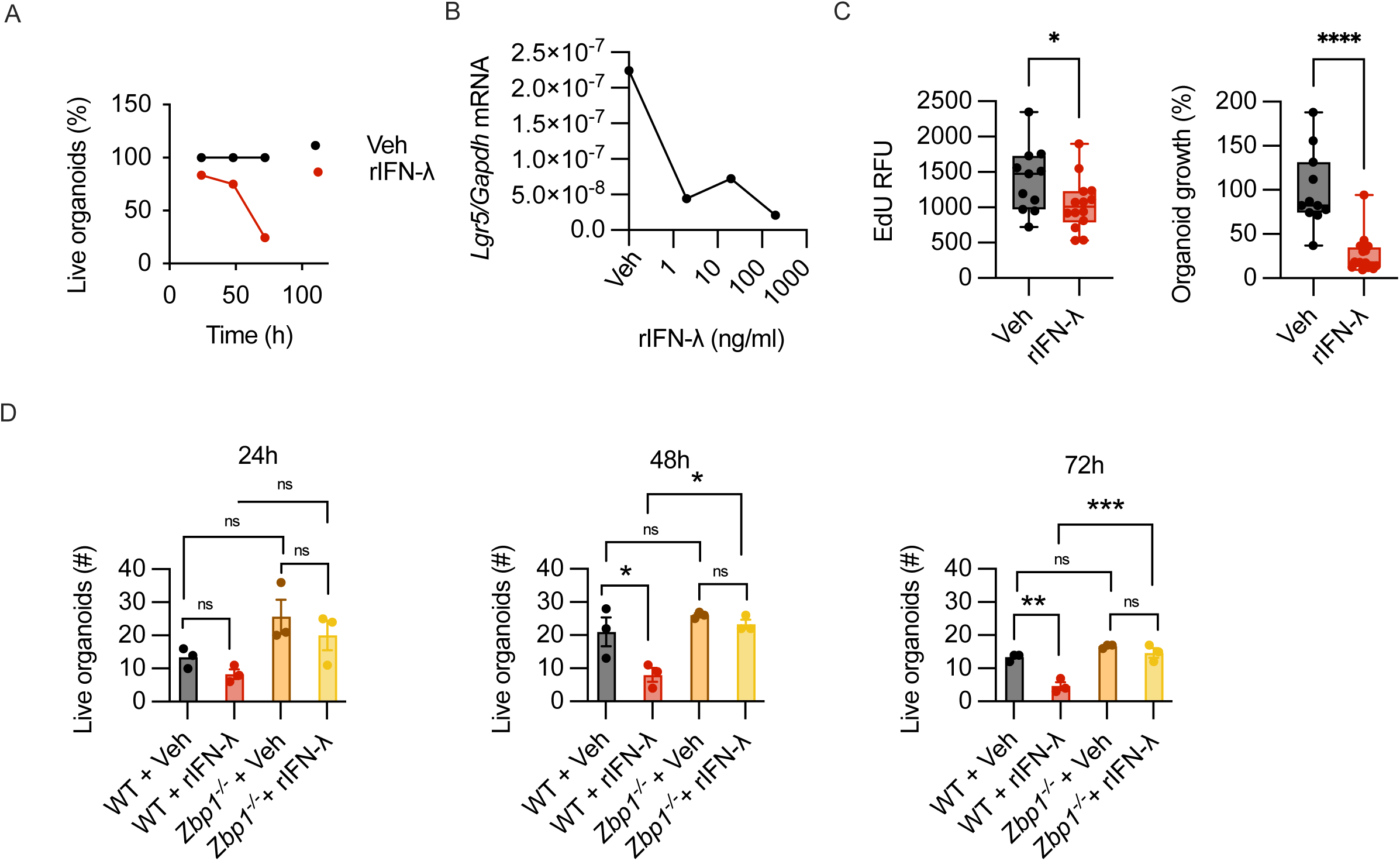
IFN-λ inhibits epithelial proliferation and survival in intestinal organoids *in vitro*. (**A**) Human duodenoids were seeded and treated (rIFN-λ), or not (Veh) with 200ng/ml of human IFN-λ2 for 72h. Cell viability was measured with CellTiter-Blue. Percentage of live organoids in rIFN-λ treated wells compared to Veh is depicted. (**B**) Mouse small intestinal organoids were grown for 6 days and then treated with mouse recombinant IFN-λ2 (rIFN-λ) at the indicated concentrations. *Lgr5* relative mRNA expression is depicted. (**C**) Human duodenoids were seeded and treated with 200ng/ml of IFN-λ2 for 72h. Organoids were pulsed with EdU for 6hr. Organoids were stained for EdU incorporation. Mean fluorescence of EdU staining (*left panel*), relative organoid growth (*right panel*), are depicted. The relative growth of organoids is measured as the % of their area over untreated control organoids. (**D**) Colon organoids derived from WT or *Zbp1^-/-^* mice were seeded and treated with 200ng/ml of rIFN-λ and imaged at 24h, 48h and 72h. The number of formed organoids is depicted. (**C**) Box plots are depicted. Each dot represents a biological replicate. Median, range and interquartile range are depicted. (**D**) Scatter box with bars are depicted. Each dot represents a biological replicate. Mean and SEM are depicted. Statistics: (**C**) Unpaired t-test. (**D**) Two-way ANOVA with Šidak correction for multiple comparisons. ns= not significant (p > 0.05); *p < 0.05;**p < 0.01; ***p < 0.001; ****p < 0.0001.

## MATERIALS AND METHODS

### Mouse strains

C57BL/6J (Jax 00664) (wild-type), *Ifnar1^-/-^*(Jax 028288), *Mrp8^CRE^* recombinase (Jax 021614), and *Vil^CRE^* recombinase (Jax 004586) mice were purchased from Jackson Labs. C57BL/6 IL-28R^−/−^ (*Ifnlr1^−^*^/^*^−^*) mice were provided by Bristol-Myers Squibb. Cells from the intestine of C57BL/6 *Zbp1*^−/−^ mice were kindly provided by Dr. A Poltorak. Cells from the intestine of B6.129S(Cg)-Stat1tm1Dlv (Stat1−/−, JAX 012606) were kindly provide by Dr. S.B. Snapper. The mutant mouse line *Ifnlr1*^tm1a(EUCOMM)Wtsi^ was provided by the Wellcome Trust Sanger Institute Mouse Genetics Project (Sanger MGP) and its funders (funding and associated primary phenotypic Information, http://www.sanger.ac.uk/mouseportal). Mice were housed under specific pathogen-free conditions at Boston Children’s Hospital, and all the procedures were approved under the Institutional Animal Care and Use Committee (IACUC) and operated under the supervision of the department of Animal Resources at Children’s Hospital (ARCH).

### Reagents and antibodies

To treat murine organoids *in vitro* and for *in vivo* administration, we used recombinant mouse IFNλ-2 (rIFN-λ) attached to polyethylene glycol (provided by Bristol-Myers Squibb), mouse recombinant IFN-β (12401-1; PBL interferonsource), anti-IFN-λ2-3 (MAB17892; R&D systems) and anti-IFNAR1 (BE00241; BioXCell), the pan caspase inhibitor Z-VAD-FMK (HY-16658B; MedChem Express), EdU (NE08701; Carbosynth). To treat human organoids *in vitro* we used recombinant human IFNλ-2 (300-02K; Peprotech). The following antibodies were used for immunoblotting: β-Actin (Mouse monoclonal; A5441; AC-15 clone; Lot# 127M4866V; Sigma-Aldrich), Rsad2 (Mouse monoclonal custom made; BioLegend) Zbp1 (Mouse monoclonal; AG-20B-0010-c100; Lot# A28231605; AdipoGen), gasderminC-2/-3 (Rabbit monoclonal; 229896; Lot# GR3317481-6; abcam), gasdermin D (Rabbit polyclonal; 20770-1-AP; Proteintech), Caspase 8 (D35G2; 4790; Lot#2; CST), Caspase 3 (9662; Lot# 19; CST), cleaved caspase 8 (Rabbit monoclonal; Asp387; D5B2; 8592; Lot#4; CST), cleaved caspase 3 (Rabbit monoclonal; D175; 5A1E; 9664; Lot#22; CST).

### DSS-colitis induction

To induce colitis, mice were given 2.5% (w/v) dextran sulfate sodium (DSS, Affymetrix) in drinking water for 7 days and were then administered water for 7 days. Where indicated in the figure legends, mice were received daily intraperitoneal injections of 50 mg kg^-1^day^-1^ rIFN-λ or rIFN-β, and, to deplete endogenous IFN-λ or to block type I IFN signaling, mice received daily intraperitoneal injections of 12.5 mg kg^-1^day^-1^ of anti-IFN-λ2-3 or anti-IFNAR1 antibody respectively. Body weight, stool consistency and the presence of blood in the stool were monitored daily. Weight change was calculated as percentage of initial weight.

### Partial body irradiation

Mice were sedated with a mix of ketamine (100 mg/ml) and xylazine (20 mg/ml) intraperitoneally. Mice then received gamma irradiation in a Best Theratronics Gammacell 40 Cesium 137-based irradiator with lead shielding of the head, thorax, and upper extremities to prevent bone marrow failure. In one sitting mice received either 11 Grey of gamma irradiation to assess tissue restitution in small intestinal crypts or 14 Grey of gamma irradiation for survival experiments. To assess proliferation, mice were intraperitoneally administered a 100 mg/kg dose of EdU in saline, final volume 500μl.

### Histological Analysis and Immunofluorescence

For morphologic analysis of irradiation experiments, fragments of the small intestine were fixed overnight in 4% paraformaldehyde (PFA) and embedded in paraffin for sectioning. Intestinal sections were stained with hematoxylin and eosin. Images were acquired with an EVOS M7000 (Thermo Fisher Scientific). Intact crypts were counted in 3 sections per animal blindly with ImageJ software. Proliferation was assessed by measuring the number of EdU^+^ cells per intestinal crypt. A minimum of 10 crypts per section and 3 sections per mouse were evaluated. For Immunofluorescence analysis, paraffin sections were deparaffinized with sequential washes in xylene and ethanol. Deparaffinized sections were then stained for EdU incorporation. All quantifications were executed in a blinded fashion.

For histology of DSS-colitis experiments, colons were flushed with PBS, flattened and rolled into a ‘Swiss roll’. Colon rolls were fixed in 10% formalin (Fisher Scientific), dehydrated in 70% Ethanol and embedded in paraffin. Paraffin sections were stained with hematoxylin and eosin and histological features were evaluated. Histological scoring was performed in a blinded fashion by assignment of a score of 1–5 to segments of the colon roll (1, presence of leukocyte infiltrate, loss of goblet cells; 2, bottom third of the crypt compromised; 3, two thirds of the crypt compromised; 4, complete crypt architecture loss; 5, complete crypt loss and lesion of the epithelial layer). Each segment was then measured with ImageJ software, and the final histological score of each sample was obtained by ‘weighting’ the score of each segment against the length of the segment and divided by the total length of the colon roll.

### EdU incorporation staining

Deparaffinized slides or organoids fixed on transwell were stained for 30 minutes with 2mM Sulfo-Cyanine5-azide (Lumiprobe) in the presence of 1mM CuSO4 and 2mg/ml Sodium ascorbate, in PBS. After EdU staining, slides were stained with DAPI (Sigma) to detect nuclei, and mounted with ProLongGold antifade reagent (Thermo Fisher Scientific).

### Crypt extraction

Small intestines were longitudinally cut and rinsed in PBS. Mucus was washed away by incubation with 1mM DTT at 4°C for 5 minutes. The tissue was moved to 10mM EDTA, 1% FBS, 1% sucrose at 37°C for 5 minutes. Samples were vortexed and small intestine fragments were moved to a new tube with 10mM EDTA, 1% FBS, 1% sucrose at 37°C for 10 minutes. Samples were vortexed. The supernatant was filtered through a 70uM strainer and kept on ice. Small intestine fragments were moved to a new tube with 10mM EDTA, 1% FBS, 1% sucrose at 37°C for 10 minutes. Samples were vortexed and the supernatant was combined with the previous fraction. The isolated crypts were resuspended in Trizol for RNA extraction and in RIPA Buffer with protease and phosphatase inhibitors for Western Blot analysis.

### Measure cytokine gene expression in the colon, small intestine crypts and organoids

Samples were collected in Trizol (Thermo Scientific) and RNA was isolated using phenol-chloroform extraction. Purified RNA was analyzed for gene expression by qPCR on a CFX384 real-time cycler (Bio-rad) using *Power* SYBR™ Green RNA-to-CT™ *1-Step* Kit (Thermo Scientific, 4389986) and pre-designed KiCqStart SYBR Green Primers (MilliporeSigma) specific for *Rsad2* (RM1_Rsad2 and FM1_Rsad2), *Lgr5* (RM1_Lgr5 and FM1_Lgr5) and IDT PrimeTime Predesigned qPCR Assays *specific for Gapdh* (Mm.PT.39a.1;).

### RNA sequencing

For targeted transcriptome sequencing, RNA (15ng) isolated from small intestinal crypts was retro-transcribed to cDNA using SuperScript VILO cDNA Synthesis Kit (11754-05; Invitrogen). Barcoded libraries were prepared using the Ion AmpliSeq Transcriptome Mouse Gene Expression Panel, Chef-Ready Kit (A36412; Thermo Scientific) as per the manufacturer’s protocol. Sequencing, read alignment, de-multiplexing, quality control and normalization was performed using an Ion S5 system (A27212; Ion Torrent). The generated count matrixes were analyzed using custom scripts in R (v 4.1.1). Differential Expression of Gene analysis was performed using the R package DEseq2 (v 1.34) with shrinkage of log2 fold changes. Volcano plots were created using the R package EnhancedVolcano (v 1.12). The differentially expressed genes with an adjusted p-value lesser that 0.1 and a log2 fold change greater than 1.5 were selected for downstream analysis. Functional enrichment analysis in Gene Ontology was performed using the R package ClusterProfiler (v 4.2) with the Biological Process terms and Benjamin-Hochberg multi-test correction with 5% of FDR threshold. Geneset Enrichment Analysis (GSEA) of hallmarks was performed using the R package fgsea (v 1.20) using the hallmark genesets (v 7.4) from the Broad Institute MSigDB or custom genesets. Leading edges of the different selected genesets were selected to build heatmaps of their expression in the different conditions and samples. The R package ComplexHeatmap (v 2.10) was used to plot the heatmaps. We used CIBERSORTx (Newman, Steen et al. 2019) to estimate the abundances of epithelial cell types using using bulk gene expression data as an input and scRNAseq signature matrices from single-cell RNA sequencing data to provide the reference gene expression profiles of pure cell populations. The scRNAseq signature matrix used to deconvolute RNAseq dataset from small intestine crypts was taken from (Haber, Biton et al. 2017). Code available upon request.

### Western blot

Western blot was performed with standard molecular biology techniques. Blots were probed for: β-Actin, Rsad2, Zbp1, gasdermin C-2/-3, gasdermin D, caspase 8, caspase 3, cleaved caspase 8 and cleaved caspase 3.

### RNAseq on IBD patient’s biopsies

The IBD biobank was generated starting with biopsy samples collected from patients suffering from CD or UC and diagnosed as clinically quiescent or in an active phase of the disease with various degrees. Controls were taken from non-inflammatory healthy portions of the colon. This investigation was registered under ClinicalTrials.gov Identifier: NCT02304666. A detailed description of the biobank as well as the RNAseq studies are described elsewhere (V. Millet et al., submitted). All raw and processed sequencing data generated in this study are accessible on the NCBI Gene Expression Omnibus (GEO) under the meta-series GSE174159: https://www.ncbi.nlm.nih.gov/geo/query/acc.cgi?acc=GSE174166 Samples were divided in three groups according to the disease status of the patient: control, inactive and active. The mean of genes for the tested hallmarks were computed by mean of their expression across the samples of each group. Linear regression analyses were performed using the geom_smooth function of the R package ggplot (v 3.3.5) with the “lm” method. Correlation analysis were performed by a Spearman test using the cor.test function of the R base package stats.

Bulk RNA-seq sequencing data was downloaded from NCBI GEO for the RISK (https://www.ncbi.nlm.nih.gov/geo/query/acc.cgi; TPM-normalized counts matrix) and PROTECT (https://www.ncbi.nlm.nih.gov/geo/query/acc.cgi?acc=GSE109142; RPKM-normalized count matrix) cohorts. For the RISK data, samples with “undetermined” histopathology data were excluded from the analysis, and IBD samples labeled as macroscopically or microscopically inflammation were categorized as “Active” with the rest as “Inactive”. For the PROTECT cohort, samples lacking histology scores were excluded from the analysis and all other IBD samples were categorized as “Active” if they had a Histology Severity Score (for chronic and active acute neutrophil inflammation) > 1 and “Inactive” if they had a Histology Severity Score of 0-1 (Boyle, Collins et al. 2017). Group comparisons between healthy controls, inactive and active IBD were performed using non-parametric t-testing (Wilcoxon test) and p values reported.

### Organoid culture and stimulation

Mouse intestinal spheroids were derived and maintained as previously described (Miyoshi and Stappenbeck 2013). Briefly, 1cm long segments of the small intestine were incubated in 5 ml of 2mM EDTA for 30 min at 4°C under rotation. Following the incubation period, the tubes were vigorously shaken, and the supernatant was passed through a 70μm strainer to collect small intestinal crypts and exclude villi fragments. The crypt compartment was collected by centrifugation, washed with advanced DMEM/F12 media (Thermo Fisher Scientific), resuspended in cold Matrigel (Corning) and plated in 40μl domes with 50% L-WRN (Wnt3, R-spondin, Noggin) supplemented medium.

Human organoids derived from healthy patients’ duodenal biopsies were kindly provided by Dr. Jay Thiagarajah. Duodenal biopsy samples were obtained from routine diagnostic endoscopy under Boston Children’s Hospital IRB protocol P00027983 and cultured with methods modified from (Sato, Stange et al. 2011). Briefly, crypts were dissociated from duodenal biopsy samples obtained from age-matched (<3 years) healthy control individuals. Isolated crypts were suspended in Matrigel and plated in 50µL domes with 50% L-WRN supplemented media.

For maintenance, organoids were liberated from the extracellular matrix by incubating in cell recovery solution (Corning) at 4°C for 30 minutes, then a single cell preparation was obtained by incubating in TrypLE express (Thermo Fisher Scientific) at 37°C for 5 minutes. Single cells were then re-plated in Matrigel with 50% L-WRN supplemented media and 10mM Rock inhibitor Y-27632. For western blot experiments, organoids were plated for 6 days. Organoids were then treated as indicated in the figure legends by adding cytokines and reagents to the medium overlaying the organoids. At the indicated timepoints, organoids were liberated from the extracellular matrix and lysed in RIPA buffer (Sigma). For microscopy of 3D organoids, freshly isolated crypts were seeded in 10μL of extracellular matrix in µ-Slide Angiogenesis (Ibidi) and overlayed with 45 μl of 50% L-WRN conditioned medium supplemented with 10mM Rock inhibitor Y-27632 for 24 hours. After 24 hours organoids were treated according to the figure legend in the presence of 1ug/ml Propidium Iodide (PI) (Sigma), and incubated in the video microscope ”Observer Z.1 Zeiss with Hamamatsu ORCA Flash 4.0 LT”, equipped with a temperature-controlled and CO2 chamber. Wells were scanned every 12 hours and mosaic brightfield and fluorescence images were taken. Organoids were identified by ImageJ and were followed over time for PI incorporation as hallmark of cell death. % of live organoids was expressed as % of organoids that never incorporated PI. Where indicated in the figure legends, organoids viability was measured with CellTiter-Blue (Promega) according to the manufacturer’s instruction. Percentage of live organoids is expressed based on relative CellTiter-Blue signal compared to untreated organoids.

For experiments with 2D organoids in Air-Liquid Interface (ALI), we followed a previously described protocol (Wang, Chiang et al. 2019). Briefly, cultured mouse small intestinal organoids were dissociated in single cells and seeded on polycarbonate transwells, with 0.4μM pores (CORNING). Initially, cells were seeded in the presence of 50% L-WRN media with 10μM Rock inhibitor Y-27632 in both the lower and the upper chamber. After 7 days, the media was removed from the upper chamber to create an ALI. Cells were maintained in these conditions for 14 days to establish a homeostatic monolayer. The ALI culture was then resubmerged with 200μL 50% L-WRN medium, for 7 days and re-exposed to air for 3 days in the presence or absence of rIFN-λ, as indicated in the figure legends. After 3 days from re-exposure to air, cells were pulsed with 10 μM EdU for 2 hours, fixed in 10% formalin and stained for EdU incorporation. Samples were examined using a Zeiss LSM 880 confocal microscope (Carl Zeiss) and data were collected with fourfold averaging at a resolution of 2100 × 2100 pixels. The percentage of EdU-positive-cells was calculated as the ratio of the number EdU-positive foci and DAPI-positive foci.

### *Gsdmc-2* and *Gsdmc-3* knock down in small intestine organoids and analysis by cytofluorimetry

GSDMC knockdown (*Gsdmc2, 3^KD^*) stable cell lines were produced using commercially designed lentivirus particles targeting mouse *Gsdmc2* (NM_001168274.1) and *Gsdmc3* (NM_183194.3) (Origene #HC108542): shRNA HC1008542A– AGTATTCAATACCTATCCCAAAGGGTTCG, HC108542B–AGTTGTGTTGTCCAGTTTCCTGTCCATGC and scrambled negative control non-effective shRNA (Origene Item no: TR30023). Lentivirus was packaged by co-transfecting shRNA and psPAX2 and pVSVG packaging plasmids into HEK293T cells. Transfection efficiency was monitored by GFP fluorescence, media was changed 24 hours post transfection and lentivirus particles were harvested in cell culture media 72 hours after transfection. Lentivirus particles were concentrated with Lenti-X^TM^ Concentrator (Takara 631232) with the manufacturers protocol. Concentrated particles were resuspended in 100% WRN conditioned media and titers were checked using Lenti-X^TM^ GoStix^TM^ Plus (Takara 631280). Only high titer lenti-particles were used to transduce duodenoids. For transduction, duodenoids were removed from Matrigel with Cell Recovery Solution and dissociated into single cells in Trypsin-EDTA for 10 minutes. Debris was removed by filtering over a 70uM Cell strainer (Stem Cell Technologies #27260) and single cells isolated at 300RCF for 10 minutes. Single cells were resuspended in 1ml of Organoid Growth Media + 500ul of concentrated lentivirus particles in a 15ml conical tube supplemented with 4ug/mL polybrene. Cells were spin-transduced in a pre-warmed 32C centrifuge in a swinging bucket rotor at 500 x g for 1 hour. Organoid pellet was resuspended in matrigel, plated onto a Corning 24 Well plate, and incubated in a 37C +5%CO2 incubator for 2 hours. After two hours 500ul of organoid growth media was added. Media was changed every other day. Transduction efficiency was assessed by GFP fluorescence, and positively transduced wells were expanded. Vehicle (Veh) treated, scramble shRNA, and *Gsdmc2, 3^KD^* duodenoids were treated with 200ng/ml of rIFN-λ in Organoid Growth Media. After 48 hours duodenoids were removed from Matrigel in Cell Recovery Solution for 1hr at 4°C, washed with PBS and resuspended in Organoid FACS Buffer (1X PBS+ 1%BSA + 2mM EDTA + 10uM Y27632). Duodenoids were stained with Zombie Aqua (Biolegend) and Calcein Red AM (Thermofisher) in Organoid FACS buffer, diluted according to manufacturer’s protocol. Cells were washed twice with PBS and mechanically disrupted by pipetting with a P200 pipet tip, then incubated with prewarmed Trypsin-EDTA for 10 minutes at 37C. After 10 minutes trypsin was quenched with Organoid FACS buffer, cells were spun at 300 x g for 10 minutes at 4°C, and resuspended in Organoid FACS Buffer for flow cytometry. Fluorescent positive gates were positioned relative to vehicle-treated and DMSO treated controls.

### Quantification and Statistical Analysis

Statistical significance was assessed by: Unpaired t-test to compare two groups, One-way ANOVA with Dunnett correction for multiple comparisons to compare 3 or more independent groups, Two-way ANOVA with Turkey or Šidak correction for multiple comparisons to compare two groups with two conditions. Kruskal-Wallis test with Dunn’s post hoc test was used to compare 3 or more independent groups when data did not meet the normality assumption. Spearman correlation analysis was used to examine the degree of association between two continuous variables. To establish the appropriate test, normal distribution and variance similarity were assessed with the D’Agostino-Pearson omnibus normality test. Statistical analyses were two-sided and performed using Prism9 (Graphpad) software or using custom scripts in R (v 4.1.1) and details are indicated in figure legends. Throughout the paper statistical significancy is defined as follows: ns, not significant (p > 0.05); *p < 0.05, **p < 0.01, ***p < 0.001, and ****p < 0.0001.

